# TATTOO-seq delineates spatial and cell type-specific regulatory programs during limb patterning

**DOI:** 10.1101/2022.03.20.482385

**Authors:** Sébastien Bastide, Elad Chomsky, Baptiste Saudemont, Yann Loe-Mie, Sandrine Schmutz, Sophie Novault, Heather Marlow, Amos Tanay, François Spitz

## Abstract

The coordinated differentiation of progenitor cells into specialized cell types and their spatial organization into distinct domains is central to embryogenesis. Here, we applied a new unbiased spatially resolved single-cell transcriptomics method to identify the genetic programs that underlie the emergence of specialized cell types during limb development and their integration in space. We uncovered combinations of transcription factors whose expression patterns are predominantly associated with cell type specification or spatial position, enabling the deconvolution of cell fate and position identity. We demonstrate that the embryonic limb undergoes a complex multi-scale re-organization upon perturbation of one of its spatial organizing centers, including the loss of specific cell populations, specific alterations in the molecular identities of other pre-existing cell states and changes in their relative spatial distribution. Altogether, our study shows how multi-dimensional single-cell and spatially resolved molecular atlases could reveal the interconnected genetic networks that regulate the intricacies of organogenesis and its reorganization upon genetic alterations.

## Introduction

The development of multicellular organisms involves the progressive specialization of cells into distinct functions according to a precise spatial and temporal blueprint. Genetically encoded programs ensure the formation of organs and other anatomical structures at the proper location through the coupling spatial identity and cell fate [1]. The ability of individual cells to acquire and interpret positional information is therefore central to the proper patterning of the embryo, and the differentiation of its multiple cell types in correctly positioned organs. Developmental abnormalities can be caused by genetic defects impacting a differentiation program *per se* [2–4], but they may also result from improper processing of spatial information [5, 6]. Furthermore, modulation of the position and activity of the different signaling centers that impose positional information during embryonic development or impairment of the ability of cells to respond to these cues have been suggested to contribute substantially to evolutionary diversity of morphologies (e.g. [7–10]). Typically, spatial position depicted as possessing an instructive role on cell fate. Yet, cell differentiation programs usually proceed in similar ways in cells exposed to different positional signals, as exemplified by the formation of muscles or bones throughout the body. This implies that spatial patterning and cell differentiation are not strictly hierarchically related but correspond to intertwined yet distinct processes.

In vertebrates, the development of the limb is a classic example of a developmental process coupling patterning and differentiation. In the mouse, forelimb buds emerge from the flank of the embryo around embryonic day E9.5 and a two-dimensional coordinate system is rapidly established by organizing centers secreting gradients of signaling molecules [11]. The proximal-distal (PD) axis is specified early by a gradient of distally secreted Apical Ectodermal Ridge (AER)-derived FGFs [12, 13] while the secretion of SHH from the posteriorly located Zone of Polarizing Activity (ZPA) defines the anterior-posterior (AP) axis [10, 14]. Within this patterned limb field, migrating somite-derived myoblasts and pools of lateral plate-derived mesenchymal cells progressively adopt different fates: muscles, cartilage, bone, tendons, and other types of connective tissue [15–17]. Although decades of genetic studies have identified the signaling pathways [12, 14, 18] and the transcription factors [19–21] engaged in limb patterning and in the differentiation of its main cell types, we still lack a comprehensive understanding on how individual cells define and adapt their cell type specific gene regulatory programs according to their position.

The recent development of single-cell transcriptomics has enabled the characterization of the diversity of cell states present in a tissue and the identification of their associated specific gene arrays [22]. Yet, the loss of spatial organization resulting from tissue dissociation still represents a major limitation for these approaches to the study of embryonic patterning. Different methods have been developed to resolve this conundrum, each with specific strengths and limitations. Unsupervised computational methods can be used to directly learn the spatial organization of the tissue from the transcriptional organization [23]. While this approach may be successful when the spatial arrangement of cell states is simple, it is not suitable for more complex systems where multiple cell states reside at a given position. Highly multiplexed single-molecule RNA-FISH methods (MER-FISH, seqFISH) [24, 25] and *in situ* sequencing methods (FISSEQ, StarMAP) [26, 27] provide direct spatial and quantitative measurements, but their detection power remains limited to a couple of thousands of genes simultaneously. Spatially resolved capture of mRNA via barcoded beads (Slide-seq, Stereo-seq [28, 29]) offers high spatial resolution for tissue sections, but in their current state, these strategies do not provide single-cell resolution and yield sparse datasets that can impede the detection of lowly expressed genes. The combination of spatial transcriptomic methods, both high-resolution imaging and laser capture microdissection, and scRNA-seq has been successfully applied in numerous contexts (e.g. [30, 31]). However, these approaches remain technically challenging, require specialized equipment and reagents and may be inconvenient to explore large samples. In the context of developmental biology, the use of known landmark genes defined by *in situ* hybridization (ISH) experiments can allow to infer *a posteriori* the location of individually sequenced cells as shown by studies in zebrafish [32] and fly [33]. However, such retrospective mapping depends on arbitrary, and sometimes hard-to-defined, detection thresholds to transform ISH signals obtained from non-linear signal amplification into binary domains of expression. Furthermore, this type of approach can only be applied when the pattern of expression of landmark genes is known and stable, which can hamper their use to analyze the consequences of mutations or other perturbations that are likely to impact the expression of the landmark genes as well. As an alternative to these methods, we developed TATTOO-seq, a simple approach which allows the characterization of thousands of cells by single-cell RNA-seq (scRNA-seq), while recording their spatial position of origin independently of their transcriptome.

### An unbiased strategy for spatially resolved scRNA-seq

TATTOO-seq is based on an optical cell-tattooing system, provided by the constitutively expressed green-to-red photoconvertible protein KikGR1 [34] (Fig. 1A). Controlling the degree of exposure to 405 nm light enables to modulate the degree of photoconversion of the pool of KikGR1 proteins, and therefore obtain distinct levels of red-to-green fluorescence ratios. This property yields an improved flexibility and resolution over simple photo-activation or photo-bleaching. Instead of simply allowing selective binary sorting (as utilized for NICHE-seq [35]), we can overlay any defined spatial pattern on the structure of interest and use the relative fluorescence intensities as a positional index to generate a more complex spatial coordinate system. Thus, as photoconverted samples are dissociated into single cell suspensions, individual cells retain their colors. They are then FACS-sorted into 384-well plates. By registering the green and red fluorescence values of each single cell as they are allocated to specific wells, we can therefore retain direct information relative to their position of origin. We combined graded photoconversion with Massively Parallel Single-cell RNA-sequencing (MARS-seq) [36], which associates a specific barcode to each individual cell to parallelly and independently determine single cell transcriptomes and positions of origin. We then used this strategy on forelimb buds from E11.5 mouse embryos (45-51 somites) and applied different photoconversion patterns with variable levels of complexity. While we were technically able to separate four degrees of photoconversion, for simplicity, we used three colors for most patterns. Under the conditions used, photoconversion was homogeneous throughout the thickness of the limb sample along the dorsal-ventral (DV). We did not detect significant color-based differences in cell viability, based on flow cytometry prior to sorting using live and dead cell markers (calcein and Sytox, respectively) (Fig. S1A). To characterize more in detail the developing limb transcriptomic landscape, we sorted ∼10,000 live cells from differently photo-converted forelimbs (embryos n = 4, limbs n = 7) and performed MARS-seq as previously described [36]. Cells with fewer than 2000 UMIs were excluded from the analysis. The remaining ∼8750 cells had a median UMI count per cell of 5980 and a median number of detected genes of 3115 (Fig. S1C). We found a high degree of correlation (R > 0.97) between samples that were exposed to different degrees of photo-conversion (Fig. S1B), indicating that photoconversion did not measurably impact gene expression. To assess the reliability of our approach in registering cell position, we compared the expression of *Meis1*, *Hoxa11* and *Hoxa13*, which are known markers of the proximal, medial and distal regions of the limb bud respectively [13], with color-patterns along the PD axis (Fig. 1B). As expected, we found a very specific overlap between cell position (as measured by color) and marker gene expression, indicating that TATTOO-seq accurately retains positional information.

**Fig. 1.**
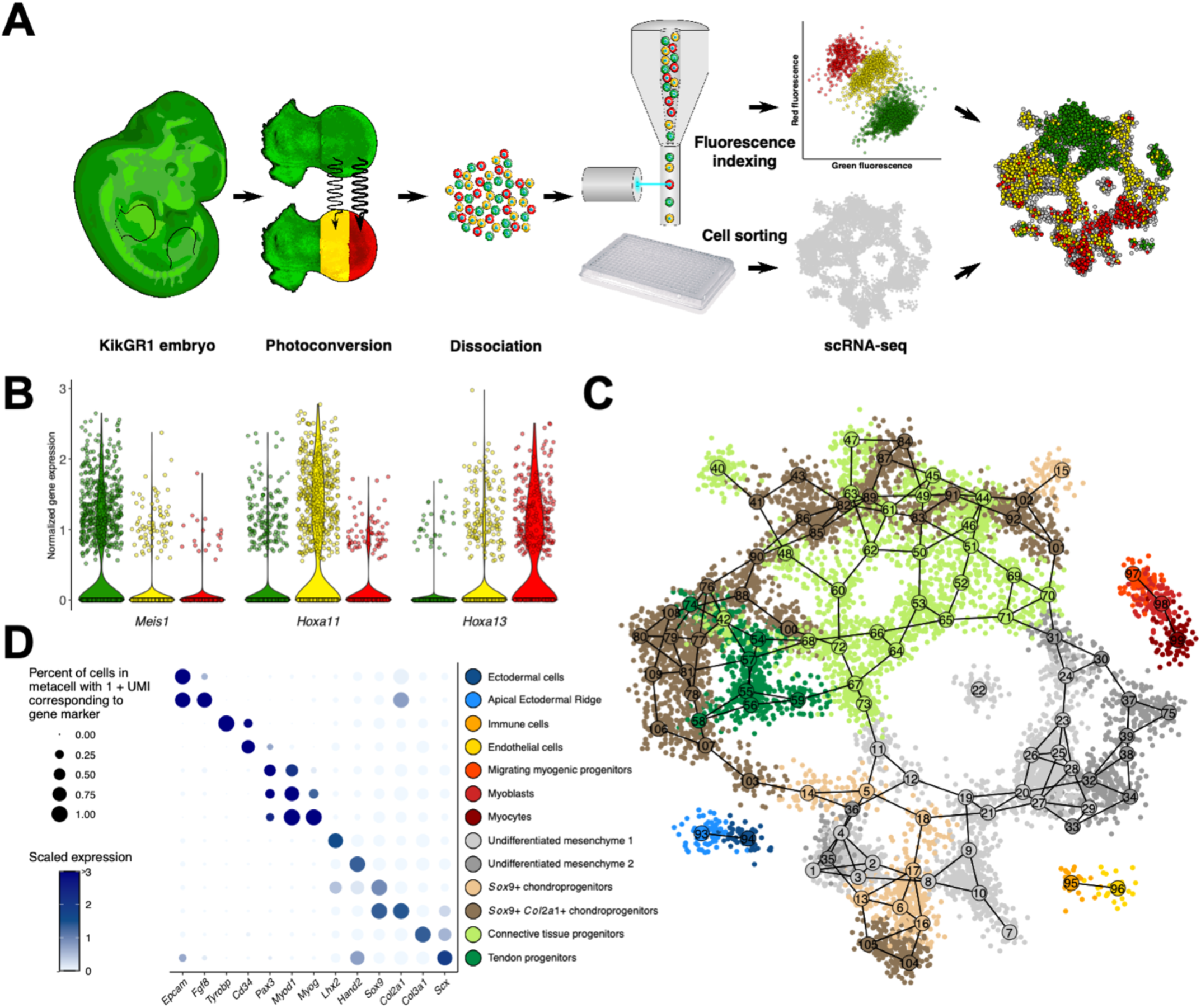
TATTOO-seq simultaneously records the position and determines the transcriptome of single-cells in the mouse limb bud. **(A)** Overview of TATTOO-seq. The structure of interest is dissected and photoconverted before dissociation. Cells are sorted into individual microwells, and their position is recorded using flow cytometry. Finally, following library preparation and sequencing, fluorescence values are re-associated to each single-cell transcriptome. **(B)** Violin plots showing the distribution of log_10_ depth-normalized counts for *Meis1*, *Hoxa11* and *Hoxa13* for each color. **(C)** 2D projection of the metacell graph. **(D)** Dot plot showing the average normalized expression of marker genes in each annotated cell type.

### A spatial molecular atlas of the developing limb bud

We used MetaCell [37] to identify transcriptionally homogeneous groups of cells and reconstructed a high-resolution transcriptional manifold model of the developing limb bud. We identified 109 metacells (Fig.1C) and assigned them to the different cell populations present in the limb bud using known cell type specific marker genes (Fig. 1D). Non-mesenchymal cell states (white blood cells, endothelial cells, and ectodermal cells) formed well-separated clusters (Fig. S2A). Ectodermal cells cluster into two distinct metacells, one corresponding to the *Fgf8*-expressing AER and the other to the dorsal and ventral ectoderms (Fig. S2B). Additionally, we detected three myogenic progenitor cell states corresponding to different stages of differentiation, from *Pax3*-expressing progenitors migrating from the somites into the limb bud, to *Myod1*-expressing early differentiating myoblasts and *Myog*-expressing differentiated myocytes [16] (Fig. S2C). As expected, mesenchymal cell states constituted the bulk of the data and formed a continuum of related cell states. The chondrogenic lineage was represented by multiple metacells, comprising condensing chondroprogenitors expressing high levels of *Sox9*, cells engaged in differentiation expressing the chondrocyte-specific collagen *Col2a1* and a few *Acan*-expressing more mature chondrocytes (Fig. S2D).

Interestingly, each of these differentiation stages of chondrogenesis are represented by multiple cell states spread throughout the mesenchymal metacell projection (Fig.1C, brown metacells). This dispersion indicates that additional components further diversify cell identity on top of the core chondrogenic program. Dense regular connective tissue progenitors (connective tissue progenitors hereafter) differentiating into tenocytes and ligament fibroblasts show expression for *Col3a1* while more differentiated tendon progenitors express *Scx* (Fig. S2E). The remaining mesenchymal cells correspond to uncommitted cells that do not express any of the canonical functional markers for chondrogenic or non-chondrogenic connective tissue. This undifferentiated mesenchyme exhibited a diversity of cell states that could be classified into two main classes based on the expression of *Lhx2* and *Hand2* (Fig. S2F). Of note, the inspection of dorsal and ventral markers (Fig. S3) suggested that, at our level of resolution, we do not capture major differences between cells positioned differentially along the DV axis. This molecular atlas revealed the diversity and complexity of the cell types and cell-states present in the limb bud. In particular, the data indicated remarkable variance in TF distribution throughout the model (Fig. S4A), showing a large number of differentially expressed TFs, even among metacells with similar cell type annotations (Fig. S4B).

### TATTOO-seq integrates multiple spatial axes into one tissue model

To delineate the spatial localization of the different cells and their associated transcriptomic states in the limb bud, we applied patterns of photoconversion (proximal-distal, anterior-posterior, distance to AER) which allowed us to capture the dimensionality of the limb as specified by its different organizing centers (Fig. 2A). The annotation of the metacell graph with this positional information immediately revealed association of the different transcriptional states to a specific location (Fig.2A). With the predictable exception of the endothelial and ectodermal metacells, the color compositions of metacells were significantly more homogeneous than expected by chance (Fig. S5). This indicated that metacells reside at precise spatial locations which supports their biological relevance as cell states. Moreover, this showed that cell position is a strong component of mesenchymal cell identity as reflected by the expression of position-specific transcriptional programs.

**Fig. 2.**
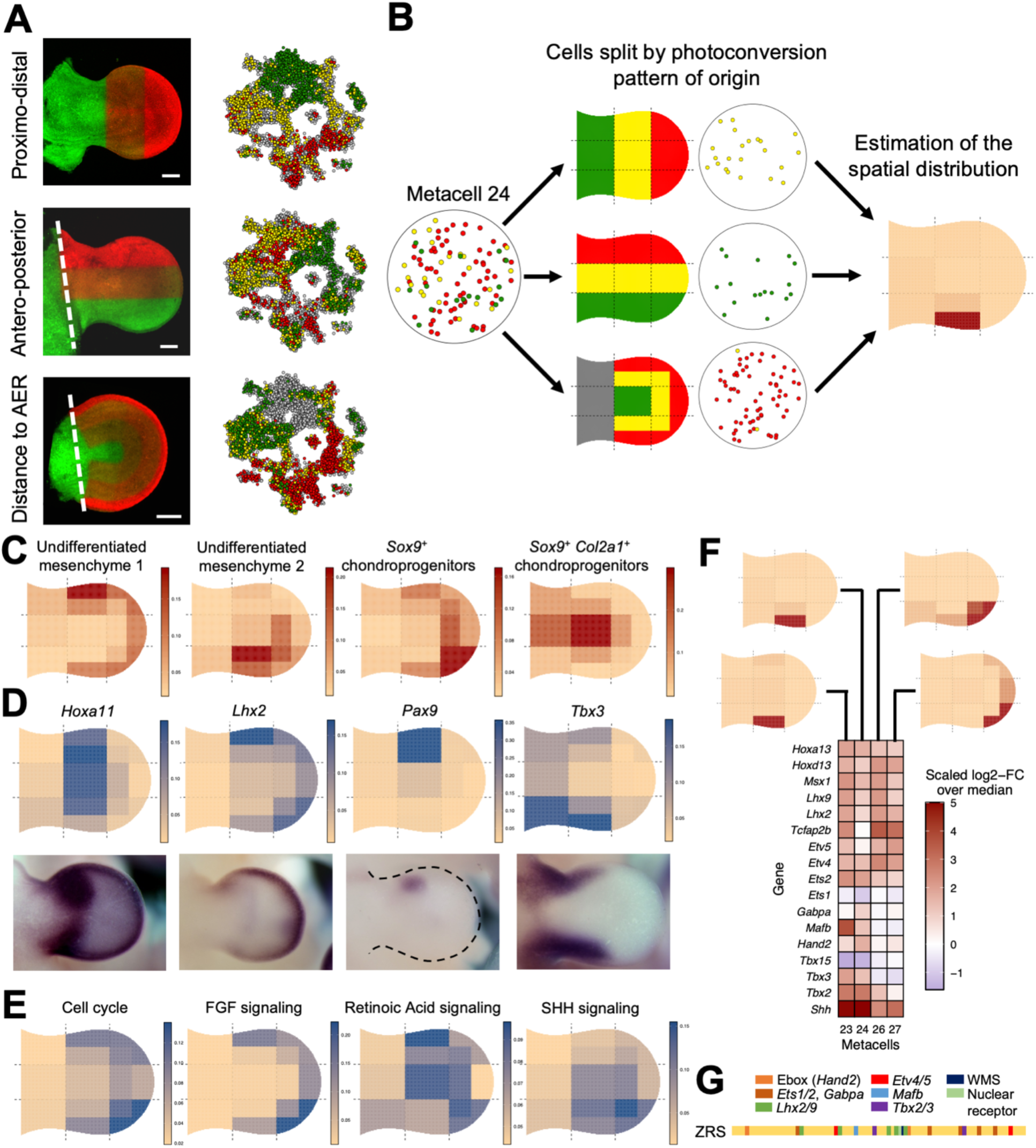
Integrating TATTOO-seq data to build a spatial map of cell states and genes expression. **(A)** Maximum intensity projection of z-stack images of each of the three photoconversion patterns: along the PD axis, along the AP axis and as a function of the distance to the AER. **(B)** Assembling the information from different patterns of photoconversion to infer the position of cell states. Three photoconversion patterns are combined into a 14-bin grid. **(C)** Spatial distribution for some aggregated mesenchymal cell types. **(D)** Virtual *in situ* hybridization (vISH) obtained *ab initio* from TATTOO-seq data. vISH accurately reproduces known ISH patterns. Images were obtained from the EMBRYS database (*38*). **(E)** Spatial projection of summarized gene expression for relevant gene sets: cell cycle genes, FGF signaling downstream targets (*Etv4/5*, *Dusp6*, *Cyp26b1*, *Spry4*, *Il17rd/Sef*), SHH downstream targets (*39*) and direct RA downstream targets (category 3 from (*40*)). **(F)** Spatial and transcriptomic dissection of the *Shh*-positive ZPA. Heatmap showing the normalized expression of TFs that have been suggested to bind to the ZRS in the four ZPA metacells. Their positions are display at the top. (**G**) Schema of the ZRS enhancer regulating *Shh* and its cohort of TF binding sites

The positional homogeneity observed within metacells across the different coordinate axes allowed us to associate each transcriptomic state coherently and confidently to its physical location in the limb bud. Even though the information provided by a single cell is limited to one axis of our coordinate system, metacells comprise cells originating from each of the three photoconversion axes. Since the abundance and location of the different cell type across the limb bud were highly similar between embryos of corresponding developmental stage, we could use this collective information to assign each metacell to a specific position in our two-dimensional coordinate system. For this, we calculated the spatial probability distribution that maximized the likelihood of observing the color composition of the metacells and position metacells into a more complex 14-bin spatial grid (Fig. 2B, see **methods**). Because our spatial model is two-dimensional, the area-normalized fraction of cells in a given bin represented an estimate of the thickness of the limb-bud at each spatial bin, assuming that the mean cell size is similar across all bins. As expected, the inferred limb bud thickness decreases as we move away from the embryo’s trunk and towards the edge of the limb bud. Transcriptionally close metacells showed markedly different spatial distribution on this 2D limb model, thereby providing an orthogonal validation of the pertinence of our metacells and demonstrates the ability of our method to infer positions of small cell populations.

### TATTOO-seq characterizes the spatial patterning of the limb bud

The possibility to assign each metacells to a specific spatial position allowed for novel ways to describe and investigate limb patterning. For instance, we determined the spatial distributions of cell types, by aggregating the spatial distributions of individual metacells assigned to the same cell type (Fig. 2C). With this approach, we showed that the two types of undifferentiated mesenchyme were located at different positions: mesenchyme 1 corresponded to cells present in the progress zone, while mesenchyme 2 cells were located in the zeugopodial posterior region consistent with their strong *Hand2* expression.

To allow the exploration of gene expression with greater precision and resolution, we developed several tools allowing to infer gene expression spatial patterns for the TATTOO-Seq spatial model. For a given gene, we can obtain a virtual *in situ* hybridization pattern (vISH) by computing the probability distribution over space for a random UMI of that gene to be found in each spatial bin. Remarkably, patterns obtained using vISH closely resembled that of the actual *in situ* hybridization experiments (Fig. 2D, Fig. S6). To facilitate exploration of limb gene expression pattern, we provide a pre-computed atlas of vISH for more than 17,000 mouse genes is available at [http://nobelmarlowlab.uchicago.edu:8888/TATTOOseq_vISH/]. We further expanded vISH to examine the spatial distribution of integrated pathways or biological process, using the geometric mean of the counts of process-specific signature genes (plus a small regularization constant) as a representation of these processes. With this approach, one can display on the limb blueprint the position of zones of cell proliferation or signaling pathways responsive-cells (Fig. 2E).

The combined spatial components of this atlas can provide further ways to visualize quantitative differences between genes across cell types and space. We can use vISHs to directly compare the expression of two or more genes (Fig. S7A) or even break down gene expression patterns by cell type or cell states. (Fig. S7B). This cell type-resolved vISH showed, for example, that the proximal anterior and distal expression domain of *Msx2* expression correspond to two vastly different cell types (undifferentiated mesenchyme and drCT progenitors, respectively) and highlight two underlying modes of regulation for *Msx2* [38]. Similarly, the expression of *Asb4* could be decomposed in anterior, central, and proximal cell type-specific contiguous domains.

Interestingly, this approach revealed diverse transcriptomic states in cell populations usually considered as homogeneous. The Zone of Polarizing Activity (ZPA), a critical signaling center which organizes limb patterning along the AP axis and influences its outgrowth by maintaining AER-derived signaling [1, 14] is located at the distal posterior margin of the limb bud and is molecularly defined by the expression of *Shh*.

We found that four different metacells expressed robustly *Shh* (23, 24, 26 and 27) and therefore represent the ZPA (Fig. 2F). While collectively the cells associated with these metacells mapped to ZPA location at the posterior distal quadrant of the limb, the different metacells showed distinct distribution, further supporting the idea that they represent actual functional domains as opposed to clustering artefacts: metacells 23 and 24 were located medially whereas metacells 26 and 27 are located more distally. Several TFs showed differential expression between these metacells, including *Hand2*, *Hox* and ETS transcription factors, which have been shown to regulate *Shh* expression [39–41], as well novel ones (Fig. 2F). We noted the presence of evolutionary conserved potential binding sites for these TFs within the *Shh* limb enhancer ZRS [7, 42, 43] (Fig. 2H), suggesting that this partition of the ZPA in distinct sub-regions may correspond to alternative ZRS-associated TFs. This hitherto hidden level of spatial and regulatory diversity in the ZPA therefore may further contribute to the diversity of limb morphologies resulting from ZRS genetic variants [44–46] as they may change *Shh* expression modulate its impact on target genes in either the proximal or distal part of the ZPA [41].

### A transcription factor code of spatial position

We next sought to use the TATTOO-seq atlas to comprehensively identify transcription factors whose expression exhibits strong spatial trends in the medial and distal mesenchymal cells. Out of 1390 TFs detected in the autopod and zeugopod mesenchymal metacells (threshold for detection: combined counts >100 UMIs), 202 showed some variable expression, ie. their expression in at least one metacell is at least 1.5-fold above its median expression across all mesenchymal metacells. We then used Spearman’s rank correlation to assess monotonic trends in gene expression as a function of the distance to the AER and along the AP axis (Fig. 3A). We uncovered a multitude of TFs whose expressions are correlated with spatial organization, highlighting the spatial heterogeneity of regulatory states in the limb bud. TFs forming anterior-to-posterior gradients included previously known genes such as *Alx4*, *Asb4*, and *Zic3* [21], but also revealed TFs with unknown functions in limb development (*Thra*, *Lmo4, Cited2*). 26 TFs formed posterior-to-anterior gradients of expression, comprising known patterning genes such as the 5’-*Hoxd* genes (*Hoxd10* to *Hoxd13, plus Evx2*) and *Hand2*. 49 TFs were expressed at higher level in mesenchymal cells close to the AER, including many of the known targets of AER-secreted FGFs and BMPs (*Lhx2/9*, *Msx1/2*, *Etv4*) and distal *Hox* genes (*Hoxd13*, *Hoxa13*). 104 genes with including genes involved in connective tissue differentiation (*Sox5/6/9*, *Scx*, *Runx1/3* [18, 47–49]) showed the opposite pattern, with weaker expression underneath the AER.

**Fig. 3.**
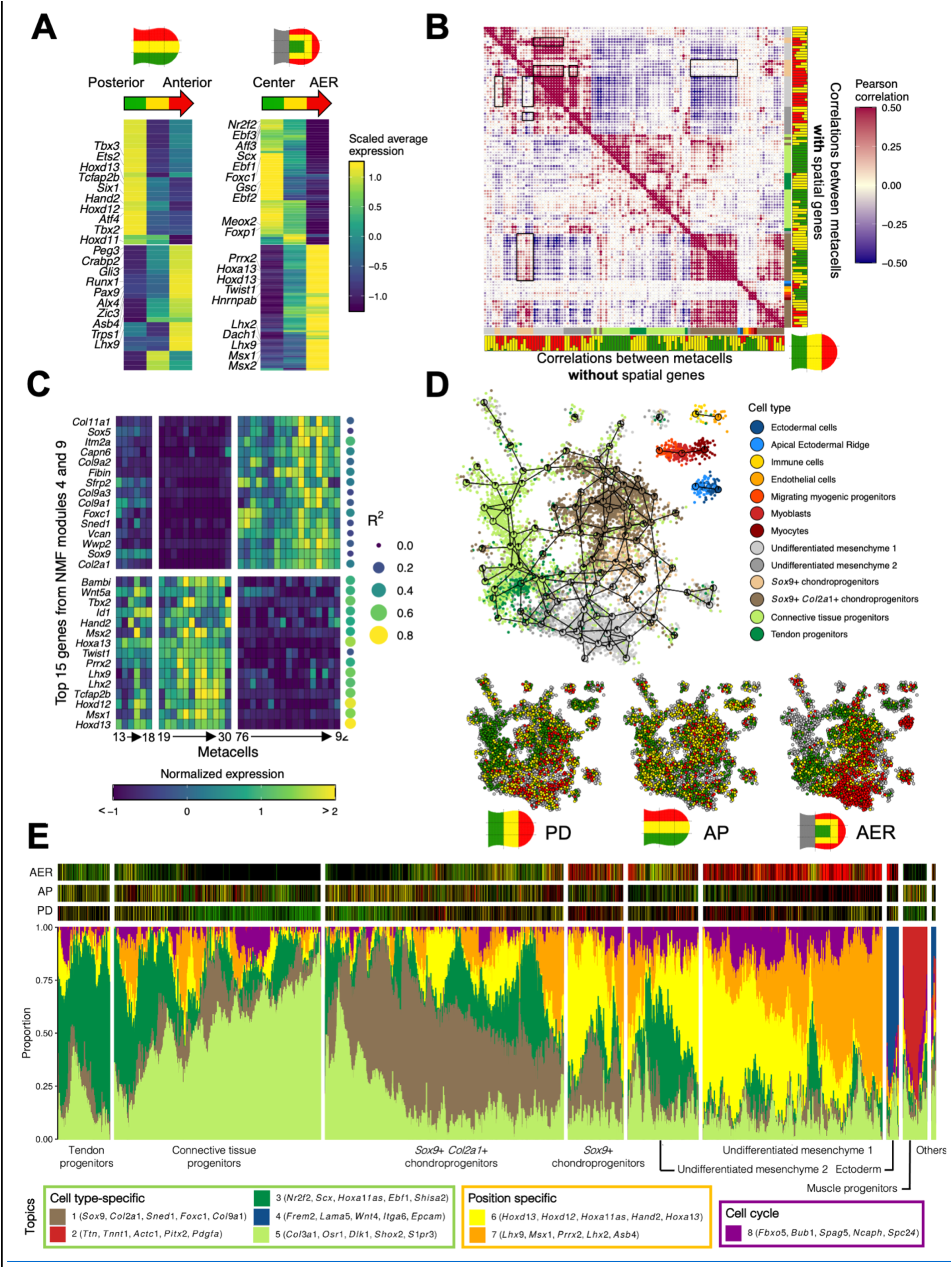
Deconvoluting positional information and cell type specific regulatory programs. **(A)** Heatmaps showing the scaled average expression per spatial compartment for TFs that exhibit spatial trends in the mesenchyme. The top 10 most significant TFs for positive and negative Spearman correlation values are annotated. **(B)** Pairwise correlation heatmap for metacells using all genes defined as HVGs (top) and excluding genes that are defined as spatial (bottom). Cell types and color composition for the PD photoconversion pattern are indicated. **(C)** Heatmap showing the normalized expression of the top 15 genes (by weight) in NMF modules 4 and 9 across metacells 13 to 30 and 76 to 92. For each gene, the R^2^ of the spatial regression is indicated. **(D)** 2D projection after clustering the TATTOO-seq dataset with no spatial genes. Cells are colored based on their previous annotation and metacells are depicted as pie charts showing the distribution of annotations. The bottom part shows cell color on the 2D projection for the three photoconversion patterns. **(E)** fastTopic structure plot showing the gradients of topic usage for each cell. The cells are grouped by cell types (endothelial and immune cells are grouped under “Others”). The legend includes the top 5 genes by estimate for each topic. Cell colors for each pattern are represented on top (black if the cell originates from a different pattern of photoconversion).

### Deconvoluting spatial patterning and cell differentiation programs

We noticed that several TFs showing spatial trends exhibited very variable expression levels within the same cell type, leading to the partition of these cell types in distinct cellular states, corresponding to distinct spatial positions. These observations could be explained by two types of underlying logics regarding how cells respond to positional cues. Firstly, the differentiation of a generic cell type (e.g chondrocytes, tenocytes) may be specified at different positions by distinct programs using different sets of TFs. Alternatively, spatial patterning genes may simply modulate the activity of an otherwise conserved core cell type-specific program. As an attempt to deconvolute the spatial and cell -type specific regulatory logics operating in the mesenchyme, we sought to classify genes as primarily carrying “spatial” or “cell type”- related information. We reasoned that genes encoding spatial information would have a similar spatial distribution of expression across cell states, while genes regulated in a strictly cell type-specific manner would show poor predictability from metacell position. We used linear regression to predict gene expression from the spatial probability distribution of each metacell for all genes and computed overall significance using F-test. After Bonferroni correction for multiple testing, we retained models for 4713 genes (out of 17537, adjusted p-value < 0.01). Strikingly, these genes include major actors downstream of signaling pathways during limb development (Table S1). We then computed and compared two pairwise correlation matrices between metacells: using all highly variable genes (HVGs), previously defined for the clustering, and using only non-“spatially regulated” HVGs, respectively (Fig. 3B). We noted two types of changes: the correlation between two groups of distal metacells decreased without spatial HVGs, while the correlation between one of these groups and proximal chondrogenic metacells increases. To better characterize this behavior, we performed Non-negative Matrix Factorization (NMF) of the gene-by-metacell expression matrix in order to identify gene modules that could explain these changes. NMF module 4 exhibits high weights for known patterning genes with strong spatial regulation (14/15 top genes with R^2^ > 0.45) while module 9 exhibits high weights for chondrogenesis genes with overall low spatial regulation (Fig. 3C). Both sets of genes include several transcription factors. The exclusion of spatially regulated genes to compute correlations decreased the influence of module 4 and therefore the similarity between metacells that simply reside at the same position and unmasked a high similarity between chondrocytes in the distal and proximal compartments. The same effect is visible by re-clustering single cells using only non-spatially regulated genes (Fig. 3D, Fig. S8). We observed that the new graph projection grouped cells together by annotated cell types, reducing the striking position-driven dispersal of the initial projection (Fig 1C). Correspondingly, cells from different positions (as shown by their color indexes) were considerably more intermixed (the original spatial signal is almost abolished for the AP axis).

To further investigate the interplay between cell position and cell fate, we used fastTopic [50, 51] to model each cell as a mixture of different (k = 8) topics which allowed to describe cell states as combinations of more or less independent modules (Fig. 3E). This uncovered a spectrum of combinatorial topic utilization consistent with the continuum of cell states highlighted by MetaCell. Notably, two anticorrelated topics are present in cells forming the undifferentiated mesenchyme 1 (topics 6 and 7). Examination of the individual cell colors shows that the relative importance of these topics followed a medial-anterior to distal-posterior axis, and the strong correlation of these topics with space. Chondrogenic progenitors are highlighted by the high prevalence of topic1 (which signature genes comprise many core chondrogenic markers), which importance did not appear to depend on the position. However, these two spatial topics 6 and 7 are still present at different levels within chondrogenic progenitors and follow the same general spatial distribution than in the undifferentiated mesenchyme. Because mesenchymal condensation and chondrogenesis are initiated as the limb develops along the PD axis, there is a strong but confounding correlation between differentiation stage and PD position. To compare chondrocyte differentiation dynamics at different positions, we therefore employed a method similar to that used to reconstruct cell trajectories across time series during zebrafish development [52], by constructing a single-cell graph of chondrogenesis in a stepwise manner, using the subspaces defined for groups of cells originating from consecutive spatial positions (instead of consecutive time points). The resulting graph was then visualized using a force-directed layout, allowing us to compare the expression of chondrogenesis genes in spite of the confounding effect of spatial compartmentalization. Interestingly, this projection reflected the heterochrony of chondrogenesis between the different spatial compartments (Fig. S9A). However, chondrogenic cells at equivalent differentiation stages did not segregate on the graph by position of origin, indicating that spatially segregated chondrocytes follow a core and largely identical differentiation trajectory, as further suggested by the dynamic of marker gene expression analyzed with ElPiGraph [53] (Fig. S9B).

These series of analyses show how a comprehensive, spatially resolved atlas can help identifying the logic and components of the core positional and differentiation programs active in a developing system. It also suggests that spatial information is not exclusively instructive, but at least in the context of chondrogenesis in the developing limb bud appear to modulate an otherwise stereotypic differentiation program.

### Integration of spatial information by position-specific regulatory landscapes

The integration of spatial information into differentiation regulatory programs can be wired at different levels. *Sox9* is a key regulator of chondrogenesis [54] with a dynamic and complex regulation during limb development [55], involving a large regulatory landscape extending over 2Mb and comprising dozens of potential limb enhancers [56, 57]. To understand how *Sox9* expression is regulated in the different limb compartments, we sought to compare chromatin accessibility at this locus along the limb PD axis. We used scATAC-seq data from wild-type E11.5 limb buds [58] to identify chromatin-accessible elements in the large regulatory domain/TAD associated with *Sox9*. We assigned a spatial position and *Sox9* activity level to scATAC-Seq cells by transferring the annotations from our TATTOO-seq clusters with Seurat/Signac [59] We then grouped cells based on position (proximal, medial and distal) and *Sox9* expression (Sox9^+^ and Sox9^-^) and conducted a differential accessibility analysis on these six groups. Despite the limited number of cells in each category (< 300 in the distal *Sox9*^+^ category), we were able to detect 6 ATAC peaks (p-value < 0.05) in the *Sox9* regulatory domain/TAD which are differentially accessible along the PD axis (Fig. S10), 5 of which correspond to previously described *Sox9* enhancers [60]. While preliminary, this data suggests that spatial information is integrated on cell type specific regulatory networks via the usage of different, position-specific enhancers.

### High-resolution characterization of cell fate and patterning alteration in mutant limbs

Patterning defects can lead to complex phenotypic consequences by perturbing cell differentiation programs as well as their spatial modulation and coordination. We sought to apply TATTOO-seq to better understand the impact of a mutation impacting the function of the AER, one of the limb’s critical signaling centers. For this purpose, we used a mouse line (DEL(Poll-SHFM)) that harbors a deletion of the enhancers that control *Fgf8* expression in the AER [61]. FGF8 is the main FGF molecule controlling the growth and patterning function of the AER, and accordingly, deletion of *Fgf8* (or the elements that control its expression in the AER) leads to severe limb malformations [12, 61]. At E11.5, DEL(Poll-SHFM) homozygous mutant forelimbs are much shorter than those of wild-type mice, display shortened bones and lack anterior skeletal elements in the forelimb zeugopod and autopod (radius and digits 1-2) as well as the stylopod (humerus) (Fig. 4A). We used TATTOO-seq as a molecular phenotyping tool to detect how the reduction of FGF signaling from the AER changes intrinsic cell states and/or their spatial distribution in the growing limb bud to lead to the observed malformations.

**Fig. 4.**
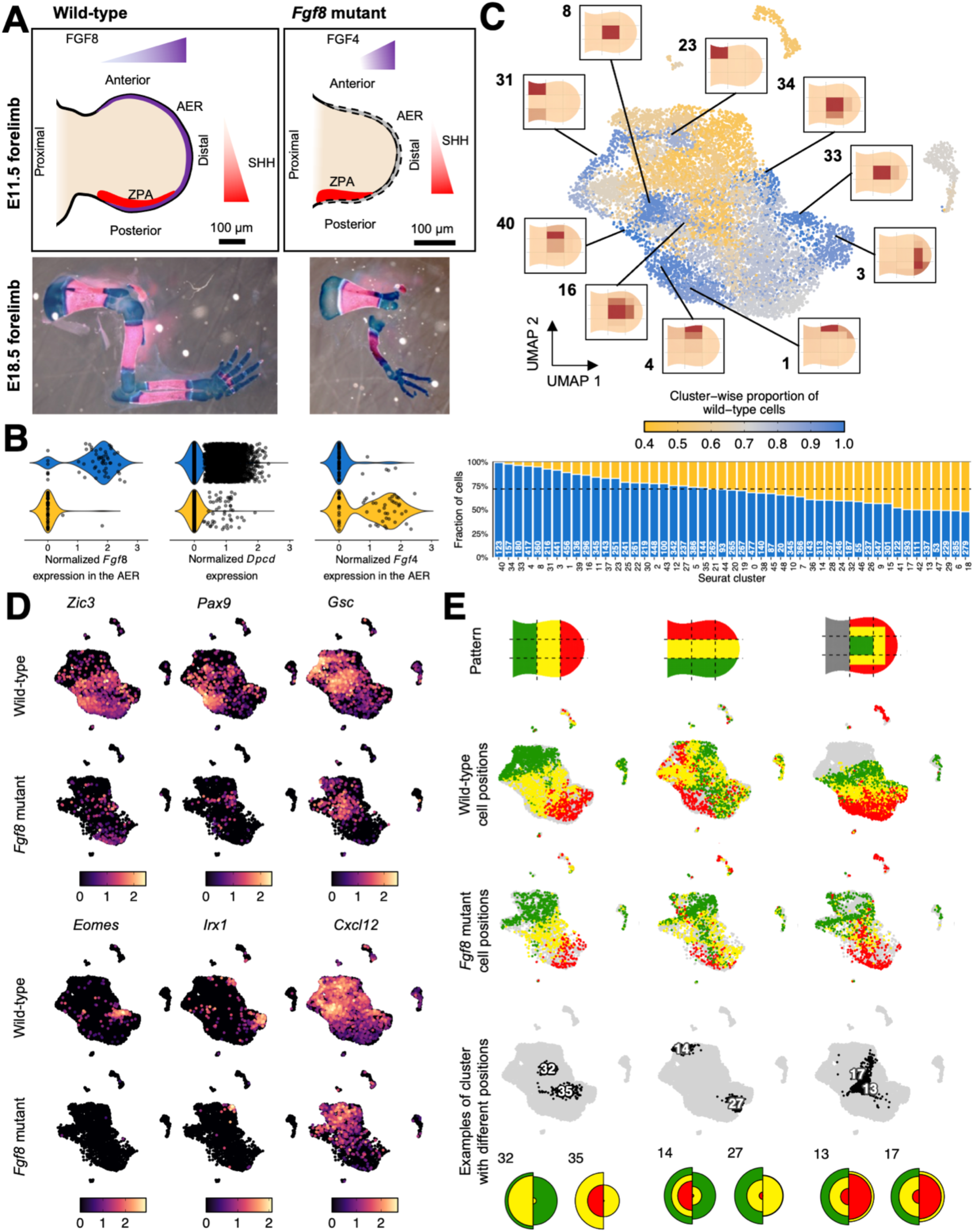
Using TATTOO-seq for high content phenotyping of a patterning mutant. **(A)** Top: schematic representation of the E11.5 wild-type and mutant forelimb buds. Bottom: skeletal preparation of wild-type and mutant forelimbs (alcian blue/alizarin red). **(B)** Violin plots showing the distribution of log_10_ depth-normalized counts for *Fgf8*, *Dpcd* and *Fgf4* for each genotype. **(C)** UMAP of the combined wild-type and mutant datasets. Cells are colored based on the fraction of wild-type cells in their cluster. The bottom part shows the fraction of wild-type cells and the total number of cells for each Seurat cluster. For clusters showing a depletion in mutant cells, a spatial projection of the wild-type cells is shown. **(D)** UMAP showing log_10_ depth-normalized counts for marker genes of the clusters showing a depletion in mutant cells. The data is split by genotype: wild-type (top row) and *Fgf8* mutant (bottom row). **(E)** UMAP showing cell colors for each photoconversion pattern split by genotype. The right part shows the location of some Seurat clusters and the distribution of colors for each genotype (left: wild-type, right: *Fgf8* mutant).

We produced a TATTOO-seq atlas for DEL(PolL-SHFM) homozygous limb buds obtained from E11.5 embryos carrying the KikGR1 transgene. Embryonic limb buds were dissected, photoconverted and dissociated as previously described for the wild-type (embryos n = 3, limbs n = 4). As mutant limbs are shorter, the AER photoconversion pattern was applied to the whole limb. After QC-based filtering, we obtained high-quality transcriptomes for 3349 mutant cells. Although this number is smaller than that of the wild-type limb, because of the size difference, it is likely that the cell type “coverage” – represented by our ability to detect transcriptionally distinct populations – is comparable. Clustering and annotation of the mutant dataset was done as for the wild-type reference. We first examined the expression of genes known to be associated directly or indirectly with the mutation and *Fgf8* in the AER. As expected, *Fgf8* expression was abolished in the ectoderm and *Dpcd*, a bystander gene included in the DEL(PolL-SHFM) deletion, was almost undetectable in the mutant cells (Fig. 4B). We could additionally observe changes reported by previous analyses of the mutant by candidate gene approaches using *in situ* hybridization, including a compensatory upregulation of *Fgf4* in the AER [12].

We then sought to characterize how the reduction of FGF signaling from the AER impacted the different limb bud cell populations. Given the spatial- and cell type-level resolution offered by TATOO-seq, we could investigate changes at multiple levels including the correct specification of the different cell types as well as changes in their transcriptomes and spatial distributions. Considering the smaller size of the mutant limbs, we first examined whether the differences present in the mutant limbs reflected a global developmental delay. For this comparison, we generated scRNAs-seq from E10.5 wild-type forelimbs (embryos n = 4, limbs n = 4; 2912 cells total after QC) and examined several molecular markers indicative of limb developmental progression in the different datasets (Fig. S11). These comparisons did not yield evidence suggesting a general heterochrony of limb development in the mutant limbs. To compare the cell populations present in the wild-type and mutant limbs, we clustered together two datasets, using a coarser clustering strategy with the Seurat package[59]. For each cluster, we computed the relative fraction of wild-type and mutant cells (Fig. 4C). Because we slightly biased the sorting strategy of the mutant cells to include more cells from the presumptive PZ (up to a 2-fold enrichment for red cells close to the AER), we could not calculate unambiguously the expected cluster-wide fractions under the null hypothesis. Yet, we found several robust differences that could not be accounted for by the sorting bias.

At this level of resolution, we did not detect any mutant-specific cell state. However, several cell types present in a normal limb bud were almost entirely absent from the mutant samples (Fig. 4C). We used the metacell clustering of the wild-type TATTOO-seq data to determine the spatial distribution of the individual cell states corresponding to Seurat clusters depleted in the mutant. Many of these depleted clusters were located in the anterior compartment of the limb, indicating notably that *Fgf8* AER deletion largely prevented the specification or growth of a spatially restricted set of cell states (Fig. 4D). The substantial depletion of these anterior cell states (characterized by the expression of *Zic3* distally, *Pax9* medially and *Gsc* proximally) provides a putative molecular basis for the absence of anterior skeletal elements at later stages. Interestingly, we also noted that posteriorly-located cell states (e.g. cluster 3), were strongly reduced in abundance in the mutant. Furthermore, the mutant exhibits a depletion of some *Cxcl12* and *Fgf18*- expressing cell states who play important roles as secondary signaling centers for connective tissue differentiation [62, 63]. While most *Sox9*-expressing cell states were shared between the wild-type and the mutant, distal chondrogenic cell states expressing *Eomes* (base of digit 4) and *Irx1* (tip of digits 2-4 [65]) were greatly reduced in the mutant (Fig. 4C and 4D). This shows that, although chondrogenesis can be initiated in an environment where FGF signaling is considerably diminished (possibly through alternative sets of *cis*-regulatory elements), spatial refinement of cell states requires genetically-encoded patterning systems. Interestingly, the depleted distal chondrogenic cells states have been proposed to act as regulation centers for TGFB superfamily-related signaling, expressing the joint-specific ligand *Gdf5* [66], the BMP-induced activin ligand (*Inhba*) [67], and the BMP antagonists *Nog* and *Chrdl1* [68, 69]. Although the depletion of these cell states is only partial, no expression of these signaling-related genes was detected in the mutant data (Fig. S12A). Absence of *Gdf5* and *Nog* signalling has been shown to lead to scapulohumeral or humero-ulnar synostosis [69], and therefore this molecular defect reported by our data could account for a missing elbow joint in DEL(Poll-Fgf8) animals.

To compare the cell types present in both the mutant and wild-type, and identify system-level and cell type-specific adjustments to altered signaling environment, we first perform genome-wide differential expression analysis using the broader cell types identified in the wild-type limb atlas, so as to maintain a sufficient power to detect gene expression changes (Fig. S12B). We found pervasive adaptation of the transcriptional states of these cell types to the reduction of *FGF* signaling from the AER with changes in expression levels of multiple genes within “conserved” cell-states (Fig. S12C and Table S2). We then compared the spatial distribution of these cell-states in the mutant and in the wild-type (Fig. 4E and Fig. S13). Only a few positional changes were detected along the PD and AER axes and most of the spatial rearrangements were observed along the AP axis. This is consistent with the lack of anterior mesenchyme cell state in the mutant leading to mutant cell states exhibiting relatively more anterior colors than their wild-type counterpart. However, we also identified opposite positional shifts along the AP axis with some cell states located more posteriorly in the mutant than in the wild-type.

## Discussion

The data and analyses provided here illustrate how the spatially resolved, single-cell level molecular signatures obtained with TATTOO-seq can offer a comprehensive and high-resolution description of the dynamic processes involved in embryonic patterning. The use of orthogonal analyses such as NMF, topic modelling and graph manifold assembly on TATTOO-seq data allowed us to appreciate the respective contributions of patterning and differentiation-specific gene modules to cell states, identify specific TFs which mediate spatial identity and constitute a first step in disentangling spatial and cell type specific gene regulatory programs. We show that positional information significantly modulates the transcriptome of limb chondrocytes, even though their differentiation appears to be controlled primarily at all positions by the same core program. Interestingly, this core differentiation program appears to operate at distinct positions through different, context-specific, modalities. These modalities associate position-specific TFs or TFs which activity is regulated by local signaling cues to distinct cis-regulatory elements which act as specific spatial integrators for common differentiation pathways. These spatial integrators are likely to subtend the molecular etiology of regionalized natural and pathological variations in morphology [70] and could be primary genetic substrate in the context of evolution for changing body shapes. The possibility to simultaneously capture cell position and identity with TATTOO-seq does not only enable to identify these elements, but provide also an integrative and multi-level description of developmental defects, as illustrated by the complex cellular, molecular and spatial changes we characterized in the mutant *Fgf8* limb buds. We expect that using such comprehensive, spatially resolved molecular fingerprint of a complex system may provide a much finer-grained, accurate representation of “mutant phenotypes”, which were so far limited to the analysis of a handful of markers and morphological features. Such approaches may better reveal how genetic perturbation altered the states of the different gene regulatory networks that control cell states and behaviors.

By independently integrating spatial position and single-cell transcriptome, TATTOO-seq constitutes a simple, robust, and flexible approach for characterizing the gene regulatory networks and cell types present during embryonic patterning and their crosstalk. Since spatial coordinates are not defined by reference-based retrospective mapping [32, 33] or pseudo-spatial ordering [23], TATTOO-seq neither requires landmark reference genes nor does it assume the homogeneity of gene expression at a given position, which are strong limitations of the aforementioned strategies. Instead of inferring individual cell positions, we labelled cells prior to dissociation based on their position, measured their position, and reconstructed cell states and their spatial distribution over the limb bud as well as gene expression patterns. TATTOO-seq shares conceptual similarities to TOMO-seq [71], though with single-cell resolution and with the simplicity and rapidity of optic labelling over physical sectioning. Because our approach relies on *in silico* cell aggregates, instead of individual cells, it is suitable for relatively sparse datasets and works without imputation. Although the spatial resolution of this study is relatively coarse (at the order of 50 to 100 µm, depending on the quadrant considered), our approach could resolve finer spatial grids as it is only limited by the precision of the photoconversion and the capacity of flow cytometry to detect graded fluorescence levels. While a genetically encoded photoconvertible protein is at the basis of our method, the recent development of photoconvertible and clickable dyes or the use of photoconvertible membrane labelling dyes [72, 73] may allow further implementation of of TATTOO-seq based approaches in non-transgenic and non-model organisms or in patient-derived samples.

## Material and methods

### Animals

Transgenic CAG-KikGR-1 mice [34] (KikGR1 hereafter, kind gift of Dr. Alexander Aulehla) were maintained inbred and genotyped either by PCR using internal primers (p1: GAAATGAAGATCGAGCTGCGTATGG and p2: CACCCTTCAGCACTCCATCACGCAC) and a standard program with 65°C annealing temperature or by assessing green fluorescence in distal phalanx biopsy. DEL(PolL-SHFM) mice [61] were maintained on an inbred C57BL/6J background.

Homozygous KikGR1 males and wild type females were crossed to generate control embryos. For mutant analysis, KikGR1/+; DEL(PolL-SHFM)/+ males and DEL(PolL-SHFM)/+ females were crossed to generate experimental and control embryos. E0.5 was defined as noon of the day the vaginal plug was detected. Embryos were collected at E11.5 and dissected in ice cold PBS supplemented with magnesium and calcium chloride. KikGR1 heterozygous embryos displayed strong and widespread green fluorescence in all tissues at all observed stages.

### Photoconversion and imaging

Samples were mounted in PBS supplemented with magnesium and calcium chloride between a glass slide and a coverslip separated by several layers of adhesive tape. Photoconversion was performed and all fluorescence images were acquired using a Zeiss inverted confocal microscope (LSM 800) using a 10X objective. Green and red fluorescence images were obtained by excitation with the 488 nm (1 mW, gain = 580, digital gain = 2) and 561 nm (1 mW, gain = 580, digital gain = 2) laser diodes, respectively. The spectra of EGFP and mCherry were used for non-photoconverted and photoconverted version of KikGR1, respectively, and adjusted such that they did not overlap. For green-to-red photoconversion of kikGR1 proteins, the 405 nm laser diode was used with variable power (100% laser power was equal to 5 mW). Full conversion (red color) was obtained after 4 iterations at 40% power while partial conversion was obtained after 4 iterations at 10% power (for the proximo-distal pattern), 9% power (for the antero-posterior pattern) or 10%/6% power (for the AER pattern). For the photoconversion along the PD and AP axes, the size of the photoconverted regions were determined to be 1/3 of the length of the limb bud (from the tip to the flank or from the anterior-most part to the posterior-most part, respectively) as measured with the ZEN software. For the AER axis, the photoconverted regions were determined to be approximately 1/4 of the radius of the dissected autopods and zeugopods.

### Tissue dissociation and fluorescence-activated cell sorting

Limb buds were dissected and incubated for 5 min in 270 µL of 0.22 Wunsch unit/mL Liberase TM (Roche, 5401119001) in EDTA-free, calcium- and magnesium-free PBS with agitation (900 rpm) at 37°C in a low-binding microcentrifuge tube (Biozym, 710176). Mechanical disruption was then performed by pipetting though a p200 gelatin-coated tip. The enzyme was inhibited by adding 30 μL of 0.5 M EDTA and volume was adjusted to 1 or 1.5 mL with calcium- and magnesium-free PBS.

1 µM Calcein Violet AM (Life Technologies, C34858) and 5 nM Sytox Red (Thermo Fisher Scientific, S34859) from Invitrogen were used as live and dead cell markers for the sorting, respectively. Excitation and emission spectra were well separated from those of the non-photoconverted and photoconverted KikGR1. Single-cell suspensions were filtered through a 35 µm BD Falcon Cell-Strainer Cap (352235) into a FACS tube which was maintained at 4°C during the experiment. Sorting was done on a BD FACSAria III with flow rate the lowest value (“1.0”). A mild spectral compensation was applied to PE-A (red) and Violet-F-450/40-A (violet) channels from the FITC-A (green) channel.

Debris and doublets were excluded from the analysis by using a sequential gating strategy relying on first SSC-A vs FSC-A followed by FSC-H vs FSC-W. Live cells were selected based on live/dead staining gating on Violet-F-450/40-A (Calcein Violet AM) versus APC-A (Sytox Red). Only Calcein-positive Sytox-negative cells were included in the analysis. Cells with a low fluorescence value in both green (FITC-A) and red (PE-A) channels were also excluded.

### Massively Parallel Single-Cell RNA-Seq

Cells were processed by MARS-seq as previously described [36]. Briefly, live single cells were sorted into 384-well capture plates containing lysis solution and barcoded poly(T) reverse-transcription (RT) primers. Unique Molecular Identifier (UMI) barcodes contain a cell-specific/well-specific label and an 8-bp random molecular tag (RMT). After evaporation of the lysis buffer, the RT reaction was performed in presence of ERCC RNA Spike-in (Ambion). Unused RT primers were digested using Exonuclease I (NEB) and wells were pooled by half plates. Second-strand cDNA synthesis was then performed, and products were *in vitro* transcribed overnight using T7 polymerase (NEB) for linear amplification. After RNA fragmentation, a partial Illumina Read1 sequencing adapter containing a plate-specific barcode was single-strand ligated using T4 RNA ligase I (NEB) and the product was reverse-transcribed. Finally, the product was purified and PCR-amplified by PCR (14 cycles) with primers containing the Illumina P5-Read1 and P7-Read2 sequences. Concentration of the purified library was assessed with a Qubit fluorometer (Life Technologies) and mean molecule size was determined with a 2200 TapeStation instrument (Agilent Technologies). Libraries were pooled and paired-end sequenced using an Illumina NextSeq 500 sequencer at a median depth of ∼45,000 reads per cell. Read1 (R1) was 70 nt long and covered the plate-specific barcode and the cDNA. Read2 (R2) was 16 nt long and covered the UMI and well-specific label.

### Quality check and read mapping

Reads were demultiplexed and count tables were built as described in [36] (scripts are available at http://compgenomics.weizmann.ac.il/tanay/). Cell-specific/well-specific labels and RMTs were extracted. Reads with low quality (Phred < 27) barcodes were discarded to prevent ambiguous or spurious assignments of reads to cells or unique molecules. R2 reads whose cell-specific or well-specific labels were unknown or mutated were discarded. R1 reads were mapped to the mouse genome (mm9) using Ensembl gene annotations (downloaded in July 2017) using Bowtie2 [74].

### Metacell analysis

We clustered the wild-type E11.5 dataset using MetaCell [37]. Briefly, an ideal metacell is a group of cells whose expression profiles are statistically equivalent to independent sampling from a single underlying transcriptional state. This is achieved by first creating a k-NN graph of the individual cells based on expression similarity and partitioning this graph into a disjoint sets of cells, metacells, which are both small (∼100 cells each) and as close to the above ideal metacell as possible.

Because plates were processed in two batches, we filtered out cells with fewer than 2000 or 2500 UMIs (lower quality transcriptomes) or more than 15000 UMIs (potential doublets). We excluded mitochondrial genes (annotated with the prefix “mt-”) from the analysis. Variable genes were selected using the parameters Tvm = 0.1, Ttot = 50 and T_top3 = 3. We then excluded cell cycles and histone genes, ribosomal genes, small nuclear riboprotein genes, some lncRNA (annotated with the suffix “Rik”) and poorly supported gene models (annotated with the prefixes “RP-”, “Gm” and “AC”) from the variable genes. In addition, we identified genes that showed a batchy expression. To do so, we performed a first clustering and, for each metacell with a sufficient number of cells from both batches (> 15), we repeatedly (10 times) randomly sampled and aggregated UMIs from 10 cells (UMIs from individual cells were downsampled to eliminate the effect of sequencing depth). If the median fold-change for a gene was > 1.6 in at least one metacell, the gene was considered batchy and blacklisted for feature selection (but not omitted from the analysis). When excluding spatial genes, we additionally blacklisted genes with adjusted p-value < 0.01 for the linear regression. A k-NN graph was built using pearson correlation and k = 100 and 500 bootstrap iterations were performed (0.75 resampling probability). Metacells were built with a minimum size of 20, K = 30 and alpha = 3. Inhomogeneous metacells were split and outlier removed (Tlfc = 3). Otherwise, default parameters were used.

### Identification of cell colors

To infer a single cell color from the intensities of the PE (red, photoconverted) and FITC (green, non-photoconverted) channels, we examined the ratio between the log of these intensities:

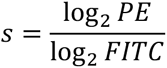

The distribution of this ratio exhibited the expected number of peaks (3 or 4) across all samples, though the exact location of the peaks and their separation varied. We therefore modeled the distribution of each individual limb separately as a mixture of skewed normal distributions whose parameters were estimated using an expectation-maximixation algorithm implementer in the R package mixsmsn [75] (“smsn.mix”, family = “Skew.normal”, nu = 3). Three components were used for the PD and AP patterns and four were used for the AER pattern. The color of each individual cell was determined using the posterior probabilities. For the AER pattern, we merged the two red-most compartments (high ratio-of-logs values) to increase the number of cells in that category.

### Modeling the TATTOO-seq photoconversion process

We treated the limb-bud as a two-dimensional shape, constructed through concatenation of Bézier curves. We then assumed that the AP and PD photoconversion patterns split this shape into three regions of equal length along the relevant axis, whose fluorescence is either mostly green, mostly yellow, or mostly red. Based on pictures of the photoconverted limb-buds, we estimated that the part of the limb-bud that was dissected for AER pattern photoconversion roughly matched the yellow and red regions of the PD pattern. We then approximated the boundaries of the AER photoconversion regions using further concatenations of Bézier curves, choosing the parameters that best fit the pictures. Note that even though the AER photoconversion split the limb-bud into four regions, we combined the two red-most regions in our model, as the number of cells collected that had red-most fluorescence was extremely low.

We then created a generative model connecting a cell’s original position within the limb-bud with its measured fluorescence. We split the modeled limb-bud’s shape into distinct spatial bins (referred to henceforth as sbins). Given the sbin containing the cell and the pattern that was used for photoconversion, it is possible to infer a distribution over the cell’s fluorescence being measured as either green, yellow or red. It is also possible to infer the probability that the cell will be completely missed by the experiment (as is the case with proximal cells when the AER photoconversion pattern is used).

Theoretically, based on the above photoconversion model, it is possible to split the limb-bud into 17 sbins which set a single, deterministic fluorescence color under all photoconversion patterns. However, using such sbins to infer cell positions led to extremely poor results, probably due to the failure of such a strictly deterministic model from handling slight deviations between the model and the experimental reality (for example, the limb-bud’s shape and the exact dissection line differed slightly between embryos). Instead, we split the limb-bud into 36 sbins based on a square 6×6 equidistant grid along the AP-PD axes. Within each such sbin the fluorescence color under the AP and PD colors is deterministically defined, while the distribution of fluorescence under the AER pattern is assumed to be proportional to the area of the sbin that is modeled to be photoconverted to green, yellow, and red respectively. We then merged together sbins whose distributions were highly similar, ending up with 14 sbins.

### Inferring cell positions

The fluorescence information provided for each cell by TATTOO-seq only limits its position to one of three or four broad spatial regions. To gain a finer spatial resolution, we utilized the reproducibility of patterning of the mouse embryo system. This allowed us to pool together spatial information collected across multiple limb buds and multiple photoconversion patterns and infer a distribution over the limb-bud’s sbins.

We will mark by λ one of the photoconversion pattern, i.e.:

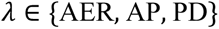

Given a subset of cells, we can calculate the regularized, empirical distribution of the measured fluorescence under each of the pattern λ:

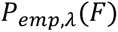

We assumed that the subset of cells defines some transcriptional criterion *M* (e.g. a cell type) which implies a distribution over the model’s sbins. Then, the sbins containing cells belonging to *M* will be distributed according to:

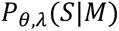

where *θ* is a parameter. Finally, the photoconversion model described in the previous section implies a distribution over the measured fluorescence given the sbin and the photoconversion patterns:

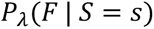

Given the above, we can calculate the expected fluorescence of cells belonging to *M*:

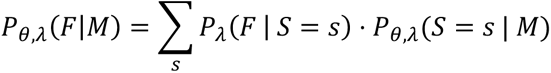

To infer the correct value of *θ*, we minimize the Kullback-Leibler divergences between the inferred distributions and the empirical ones:

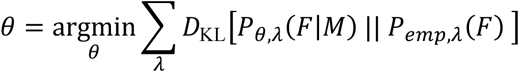

As the distributions *P*_θ,λ_(*S*|*M*) are discrete, the parameter *θ* can be taken to be the complete definition of the distribution. To solve the minimization problem, we used a constrained interior-point algorithm, implemented in the scipy.optimize.minimize() function (from the scipy python package) with a method parameter of “trust-constr”. To ensure convergence to a proper distribution function, the sum of the probabilities was constrained to 1:

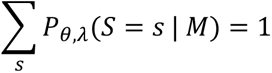

And the value of the probability of each sbin was constrained to a segment which strictly contain the [0, 1] segment:

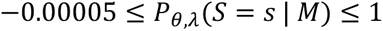

When calculating the Kullback-Leibler divergences, the individual sbin probabilities were clamped to the [0.001, 1] segment. Following the convergence of the algorithm, any negative sbin probabilities were set to 0, and the probabilities renormalized to ensure they sum to 1.

### Inferring positions of metacells

The cells of a single metacell form a natural candidate for applying the cell positioning algorithm described in the previous section. We expect that the expression profiles of most of the limb-bud’s cell types depend on spatial signaling fields. We can therefore assume that cells with a highly similar expression pattern (as the cells of a single metacell are guaranteed to be) are also tightly localized within the limb-bud. The use of a metacell also provides a natural definition to the transcriptional criterion *M*, i.e. the inferred sbin distribution describes the expected spatial localization of new cells whose expression profile is close to the metacell’s centroid.

### Metacell specific transcription factors

The metacell-specificity of transcription factors was estimated using a method similar to the “roc’’ method in Seurat. For each gene, the empirical CDF was computed based on the rescaled (from 0 to 1) gene expression in metacells. The area under the curve (AUC) was then computed. Genes that have very specific patterns of expression (i.e., expression in very few metacells) display large values of AUC because most metacells exhibit low values of gene expression.

### Chondrogenic trajectory and pseudotime

Each chondrogenic progenitor was assigned to a compartment along the PD axis based on its color. Cells originating from other photoconversion patterns, were assigned to the compartment for which their metacells had the highest probability. We performed PCA on the top 2000 variable genes within each compartment (thus excluding variation resulting from spatial compartment-specific genes) and built graphs connecting each cell to its 40 nearest neighbors within its compartment. Projecting medial cells into the distal PC space, we built a graph between distal and medial cells. Similarly, we built a graph connecting medial and proximal cells in the medial PC space. Edges were then filtered to keep at most 10 mutual edges. The resulting graph was projected using the DrL graph layout. ElPigraph [53] was then used to compute the pseudotime for each cell. Gene expression as a function of the pseudotime was computed and smoothed using loess regression.

## Acknowledgements

We thank Dr. Fujimoto and the RIKEN Laboratory for Animal Resources and Genetic Engineering for given us access to the KikGR-1 transgenic line (ref. CDB0201T-1), and Dr. Alexander Aulehla (EMBL) for sharing with us these mice. We acknowledge greatly the contribution of the members of the animal facilities at the Institut Pasteur. We thank members of the Spitz, Marlow, and Tanay labs for sharing reagents, ideas, and comments. We also thank Dr. Chloé Moreau for helping with the pilot photoconversion experiments. We particularly acknowledge the contribution of Dr. Elodie Brient-Litzler and Dr. Jean-Christophe Olivo-Marin (Institut Pasteur, CiTech) in the development of a single-cell initiative and technological platform at the Institut Pasteur, which was essential to this project. This project was also directly supported by the Institut Pasteur, the Region Ile-de-France (program Sesame) and la Fondation pour la Recherche Medicale (programme Equipe 2018 to F.S.).

---

**Fig. S1.**
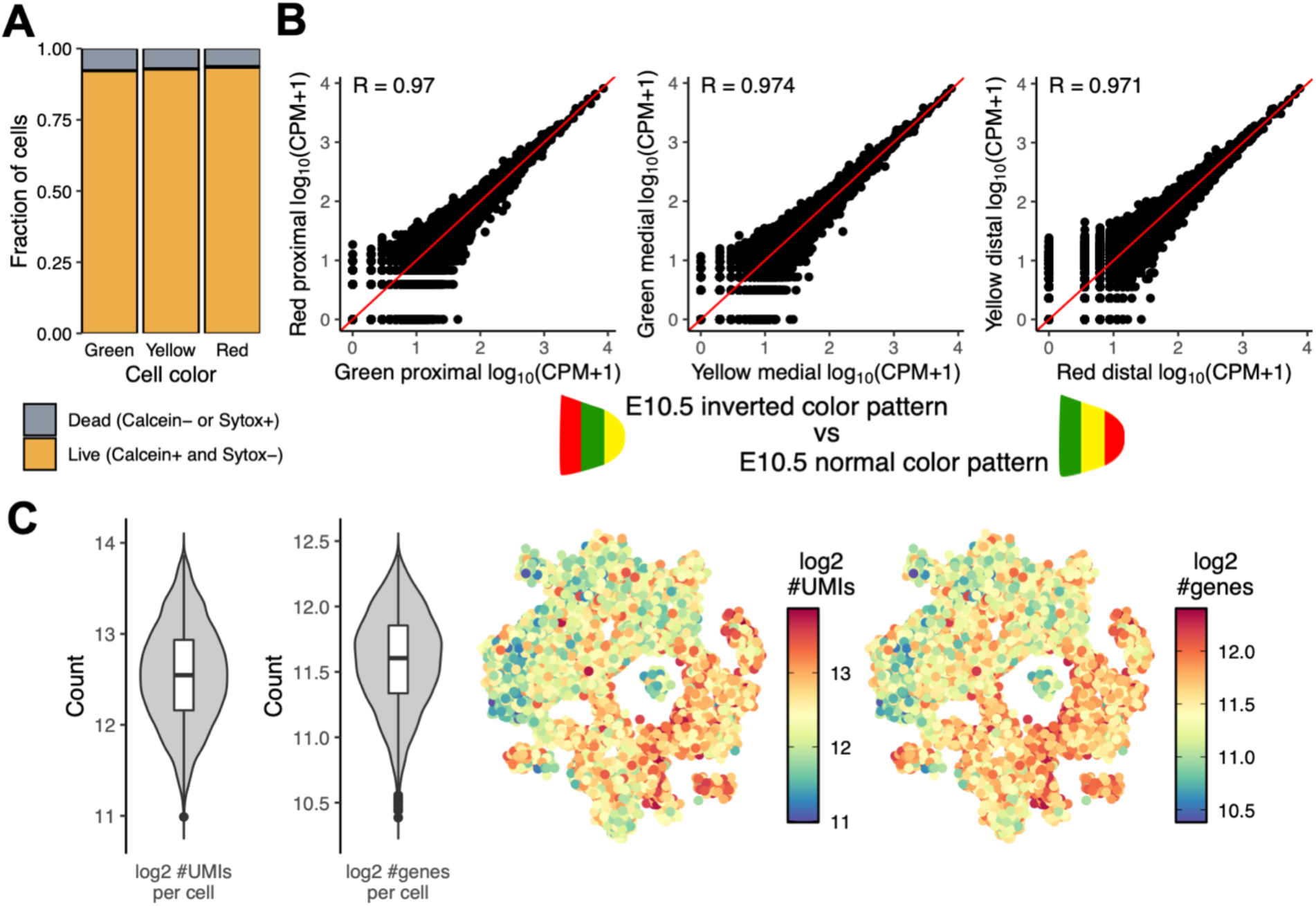
Assessing the impact of the TATTOO-seq method on cell viability and transcriptome and checking the quality of the data. **(A)** Fraction of live and dead cells (using Calcein and Sytox) assessed by flow cytometry for each color. The fraction of dead cells is independent from the color. **(B)** Comparison of aggregated gene expression for different degrees of photoconversion on the same spatial compartment. **(C)** Violin and 2D projection showing the distribution of the number of genes and number of UMIs detected per cell. The number of detected genes and UMIs is slightly higher in distal cells which are more actively proliferating.

**Fig. S2.**
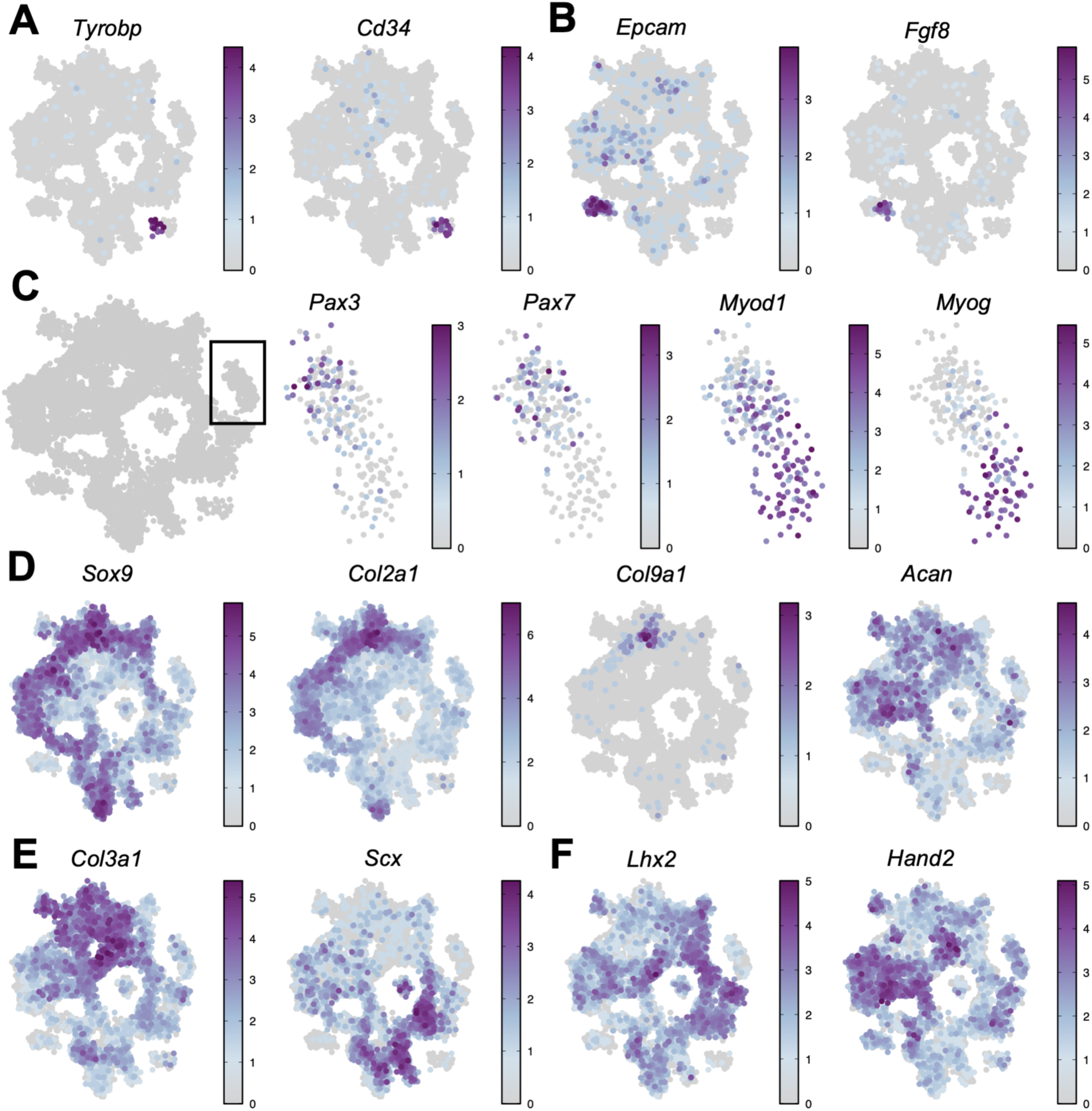
Expression of cell type specific marker genes. 2D projection of the metacell graph. Cell color represents log_10_ depth-normalized counts for marker genes of **(A)** immune and endothelial cells, **(B)** epithelial cells and AER cells, **(C)** stages of muscle progenitor differentiation, **(D)** chondrogenic progenitors, **(E)** dense regular connective tissue progenitors and **(F)** different populations of undifferentiated mesenchyme.

**Fig. S3.**
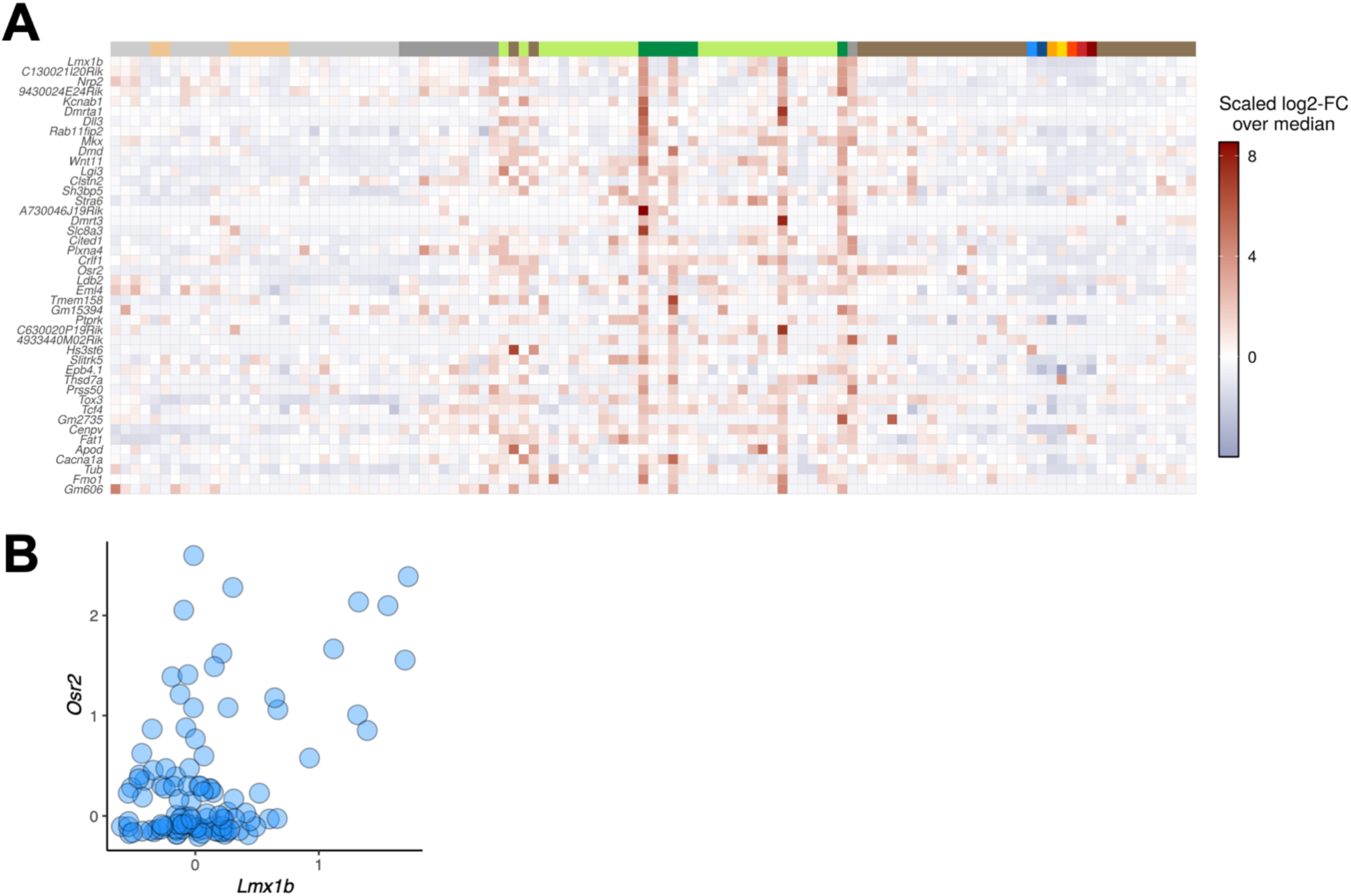
DV organization of the limb cell states. **(A)** *Lmx1b* was used as a marker of the dorsal mesenchyme [76]. Heatmap showing the normalized expression of genes whose expression is loosely correlated or anti-correlated to that of *Lmx1b* (|R| > 0.4). **(B)** Scatter plot showing the expression of *Lmx1b* and *Osr2* for all metacells. Metacells exhibiting high levels of *Lmx1b* showed high levels of *Osr2*, a key regulator in connective tissue differentiation, suggesting that connective-tissue differentiation at this stage is preferentially occurring dorsally, as suggested by *Osr2 in situ* experiments [77].

**Fig. S4.**
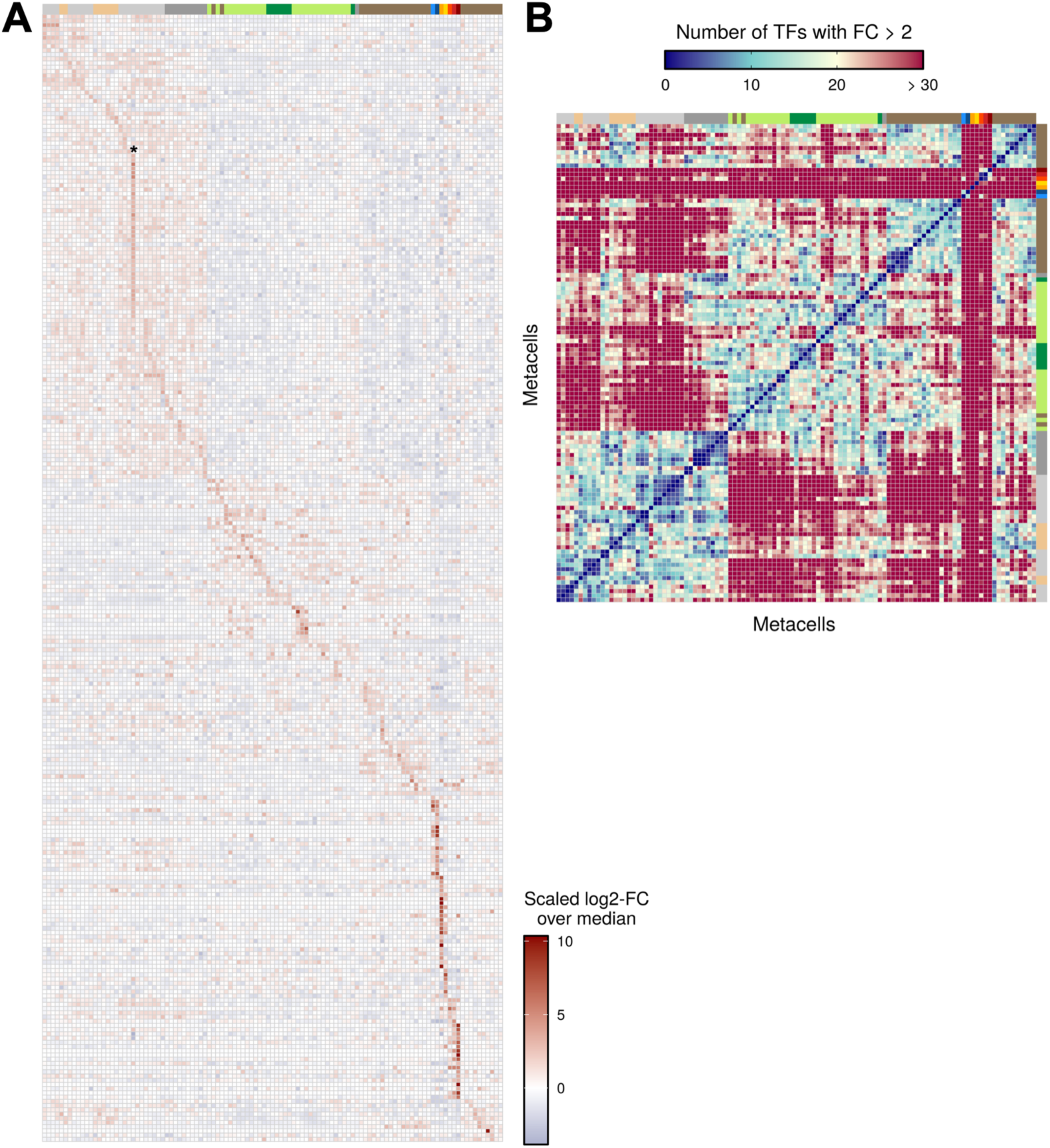
Metacells exhibit a hierarchical and specific combinatorial code of TFs. **(A)** Heatmap showing the normalized expression of genes with loosely specific gene expression (AUC > 0.5) across all metacells. * marks metacell 22 which groups cells of overall lower quality. **(B)** Heatmap showing the pairwise number of TFs that show different expression levels (FC > 2). Cell types are annotated using the same color code as in **Fig. 1**.

**Fig. S5.**
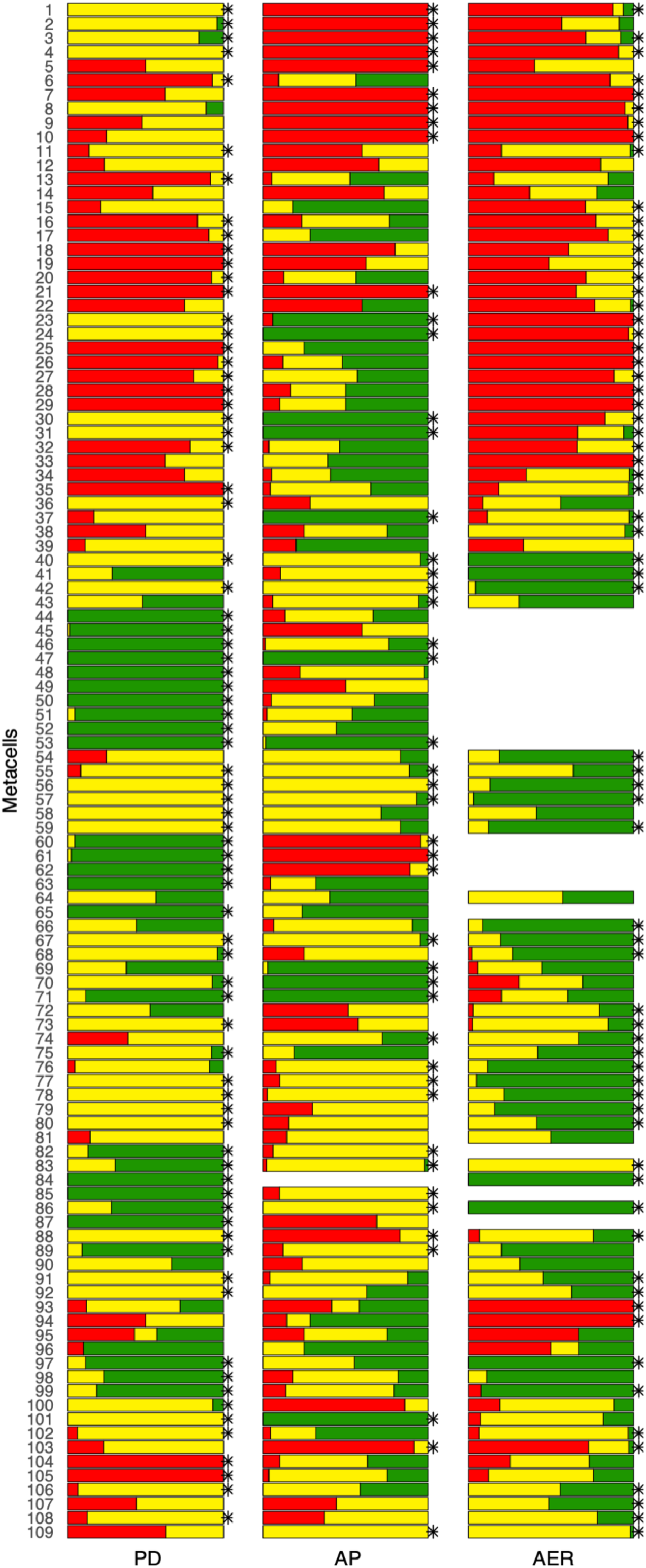
Metacells are spatially homogeneous. We employed Shannon entropy as a metric for measuring color heterogeneity. Monte Carlo simulations (n = 100000) were used to test for positional homogeneity. Stacked bar plots showing the distribution of color for each metacell in all three photoconversion patterns (PD, AP and AER). Clusters labeled with a start (*) display significantly lower Shannon entropy than expected by chance (Bonferroni corrected p-value ≤ 0.01).

**Fig. S6.**
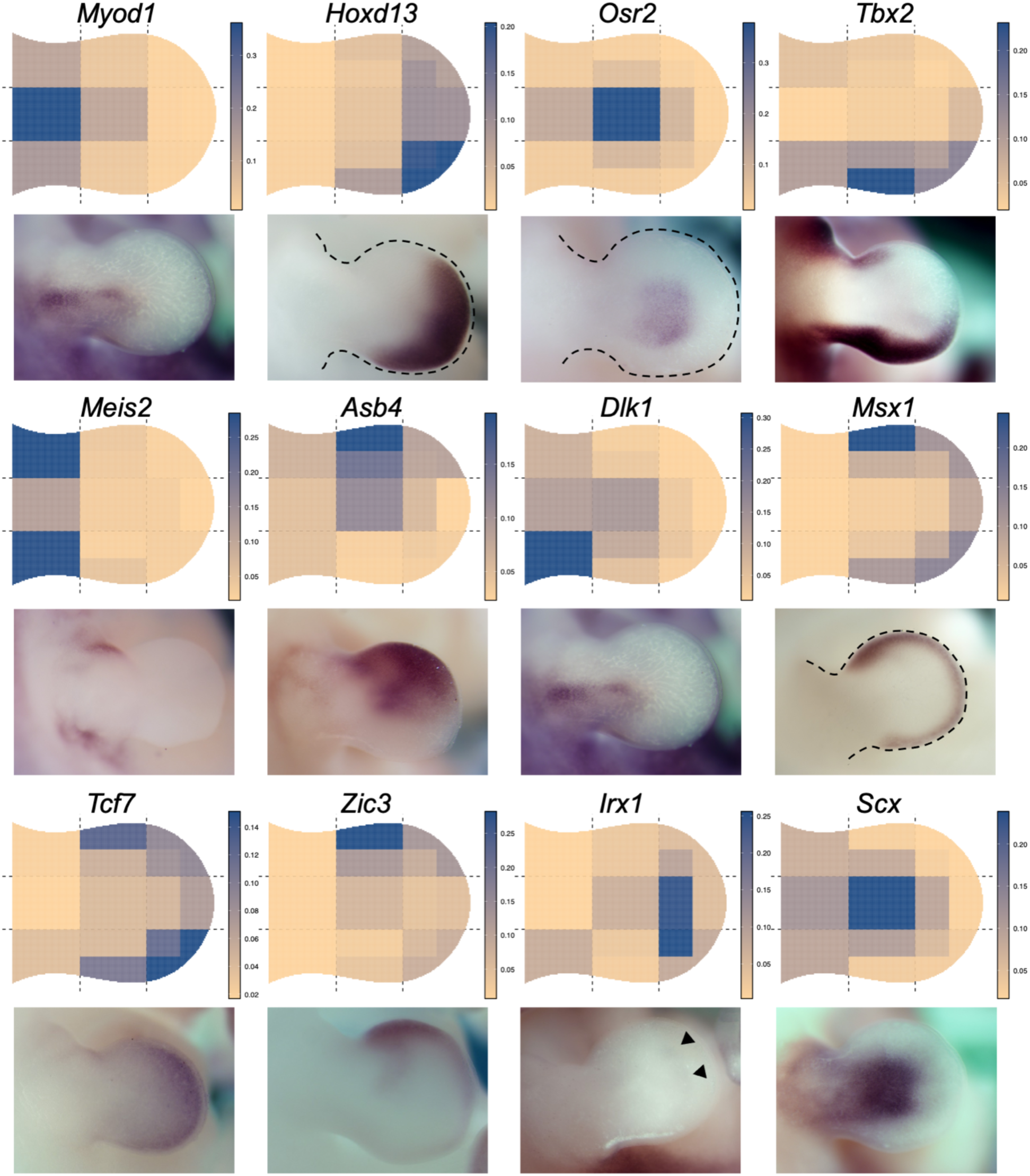
vISH match the gene expression patterns obtained by classical *in situ* hybridization. vISH and the corresponding classical ISH patterns for twelve TFs involved in limb development. ISH images were obtained from the EMBRYS database [78, 79].

**Fig. S7.**
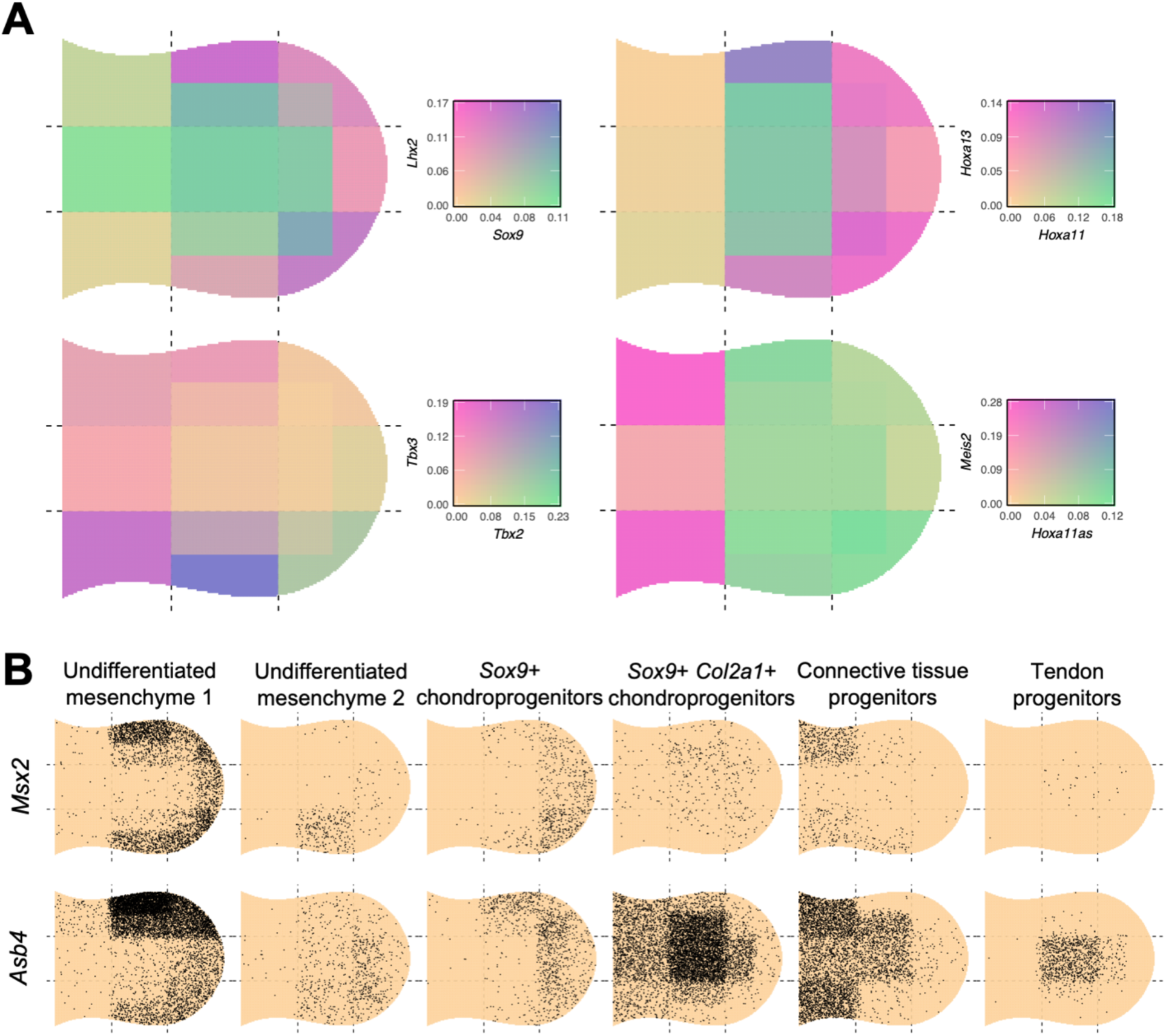
Double virtual *in situ* hybridization (vISH) and cell type-specific vISH. **(A)** Double vISH uses a bivariate color scale to represent the spatial overlap of expression patterns. **(B)** vISH can be deconvolved into cell type-specific vISH. To take into account the different dynamic ranges across cell types, up to 10,000 UMIs are sampled according to the expected number of counts for each spatial bin.

**Fig. S8.**
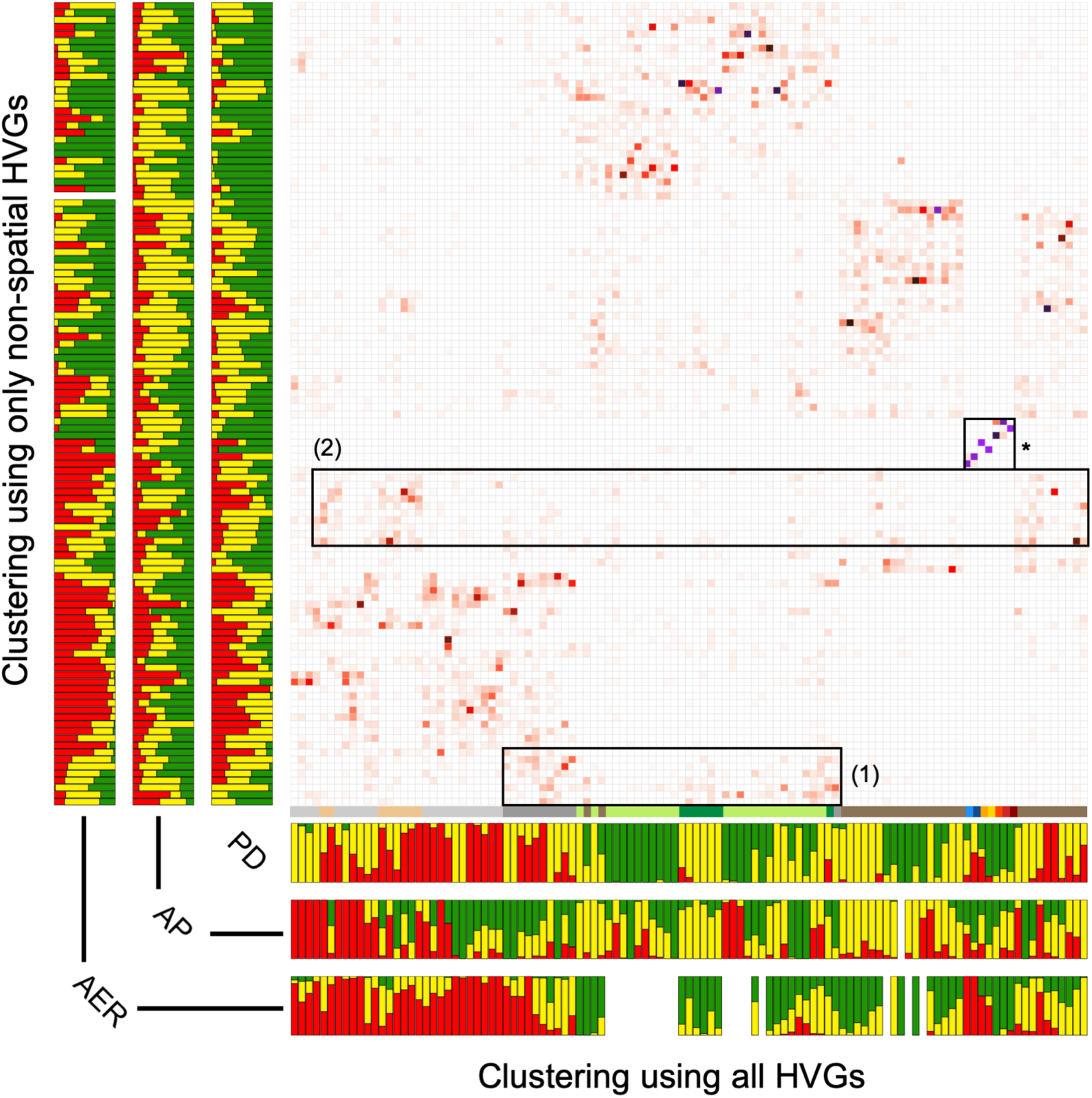
Confusion matrix between the two dataset clusterings. Confusion matrix representing the redistribution of cells between the clustering using all the highly variable genes and the clustering using only non-spatial highly variable genes. Darker values (black, purple) indicate high correspondence between metacells of the two clusterings. The color distribution is indicated on the side of the heatmap. Non-mesenchymal clusters are unaffected (*) while some mesenchymal clusters are split and merged. Some no-HVGs metacells clusters receive contribution from type 2 undifferentiated mesenchyme and dense regular connective tissue progenitors metacells (1), and from distal chondrogenic progenitors and proximal *Sox9*^+^ *Col2a1*^+^ chondroprogenitors (2). Blacklisting of spatial HVGs additionally results in extensive cell type internal reorganization. Abrogating spatial information potentially leads to artefactual clustering if cell types are homogeneous.

**Fig. S9.**
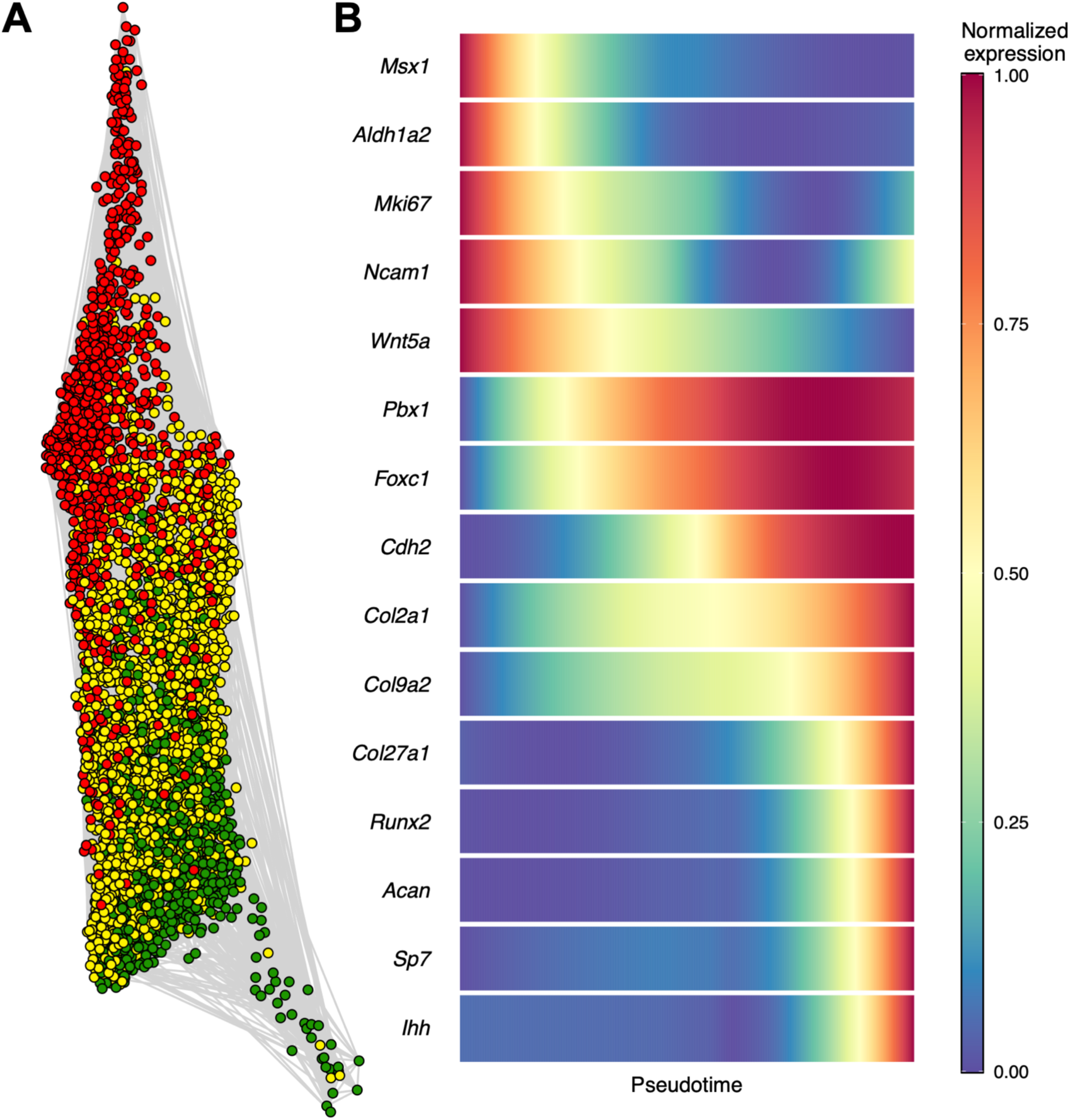
Eliminating confounder effects of cell position reveals overlapping chondrogenic transcriptomic states. **(A)** 2D projection of the k-NN (k = 10) graph of chondrocytes using the Distributed Recursive (Graph) Layout algorithm. The procedure used to generate the graph is described in the **methods** section. **(B)** Pseudotime was assessed using ElPiGraph and gene expression as a function of pseudotime was estimated and smoothed using a loess regression for various stage-specific chondrogenesis markers.

**Fig. S10.**
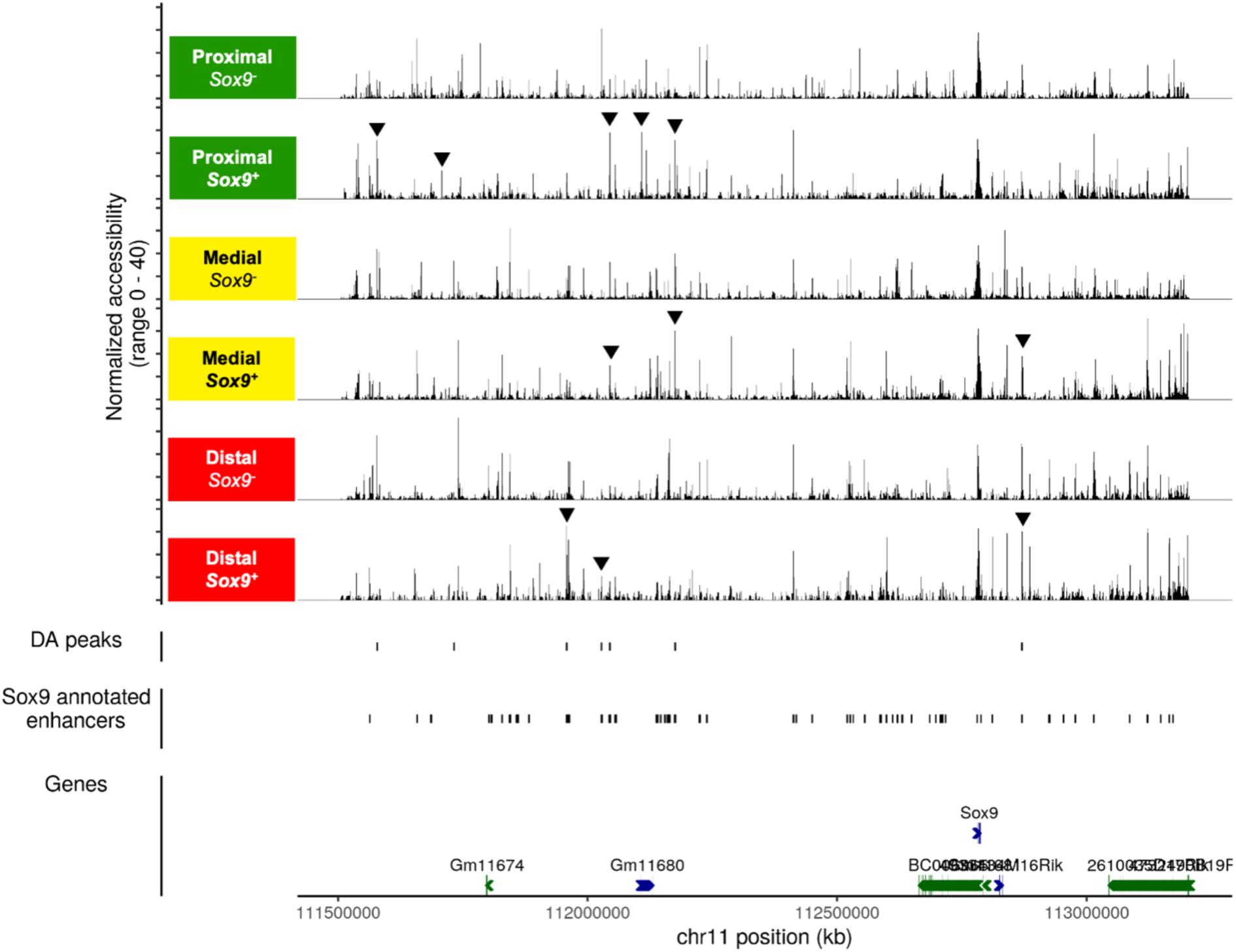
Transferred cell identity and position from TATTOO-seq to scATAC-seq points to spatially regulated accessibility landscape at the *Sox9* locus. Cell identity and position was transferred from our TATTOO-seq data to limb scATAC-seq data using Seurat’s TransferData function. Accessibility profiles were aggregated by transferred position and *Sox9* expression levels and displayed at the top. All peaks in the *Sox9* regulatory domain were tested for differential accessibility. Differentially accessible (DA) peaks are indicated as well as previously described *Sox9* enhancers [60].

**Fig. S11.**
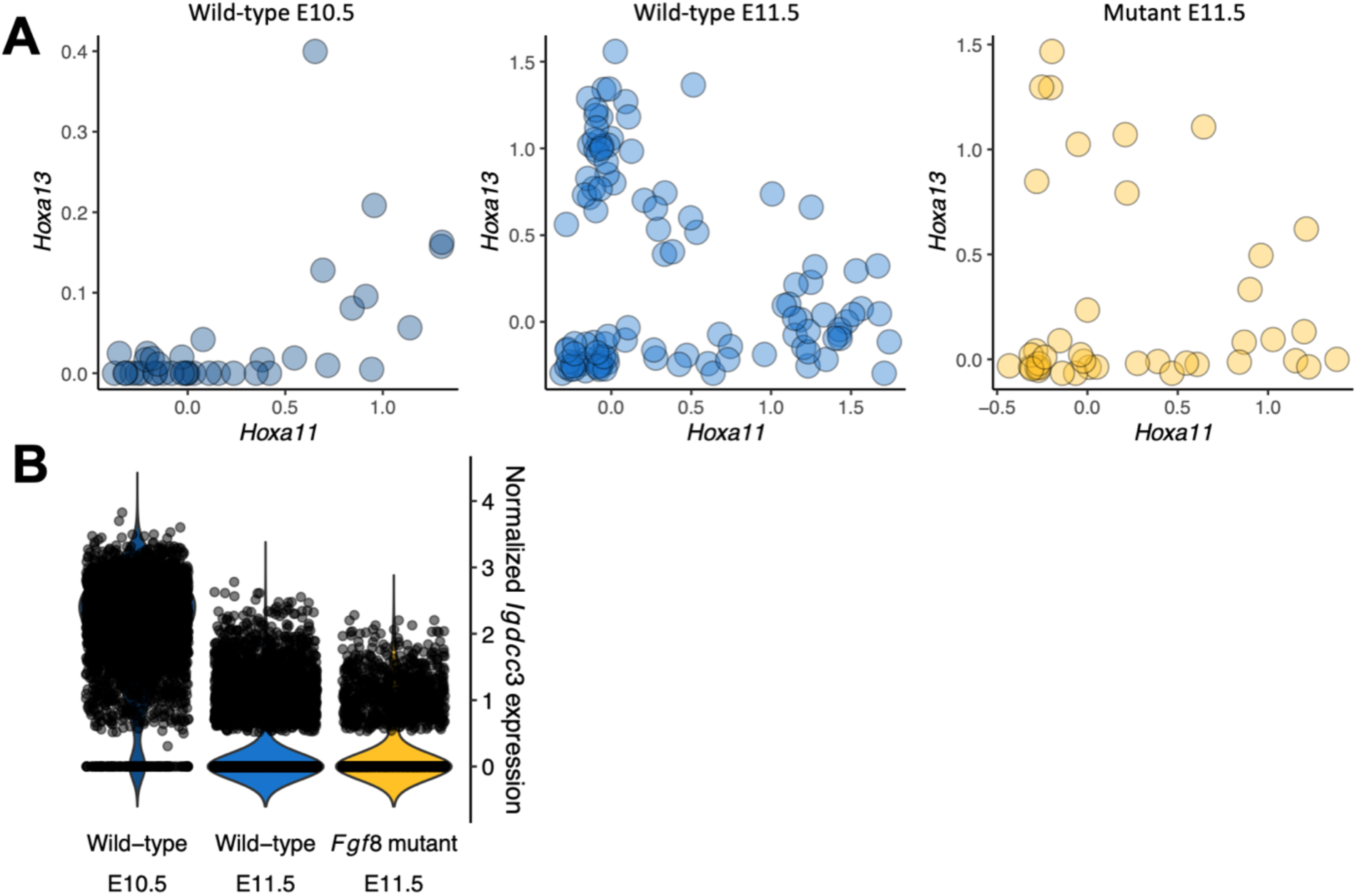
Differences between wild-type and *Fgf8* mutant cells do not reflect a global developmental delay in mutant limbs. **(A)** Scatter plot showing the normalized expression of *Hoxa13* and *Hoxa11* in E10.5 and E11.5 wild-type metacells and E11.5 mutant metacells. In the mutant limbs, *Hoxa13* and *Hoxa11* were expressed in distinct cell-states, corresponding to the largely mutually exclusive territories observed in wild-type E11.5 limbs, and contrasting with the nested expression found earlier at E10.5 [80, 81]. **(B)** Violin plot showing the log_10_ depth-normalized expression of *Igdcc3* in E10.5 and E11.5 wild-type cells and E11.5 mutant cells. *Igdcc3* has been reported to be a sensitive temporal marker of limb development, showing strong expression at E10.5 but minimal expression at E11.5 . We found its expression greatly reduced in both wild-type and mutant E11.5 scRNA-seq data, while it is robustly detected in E10.5 samples.

**Fig. S12.**
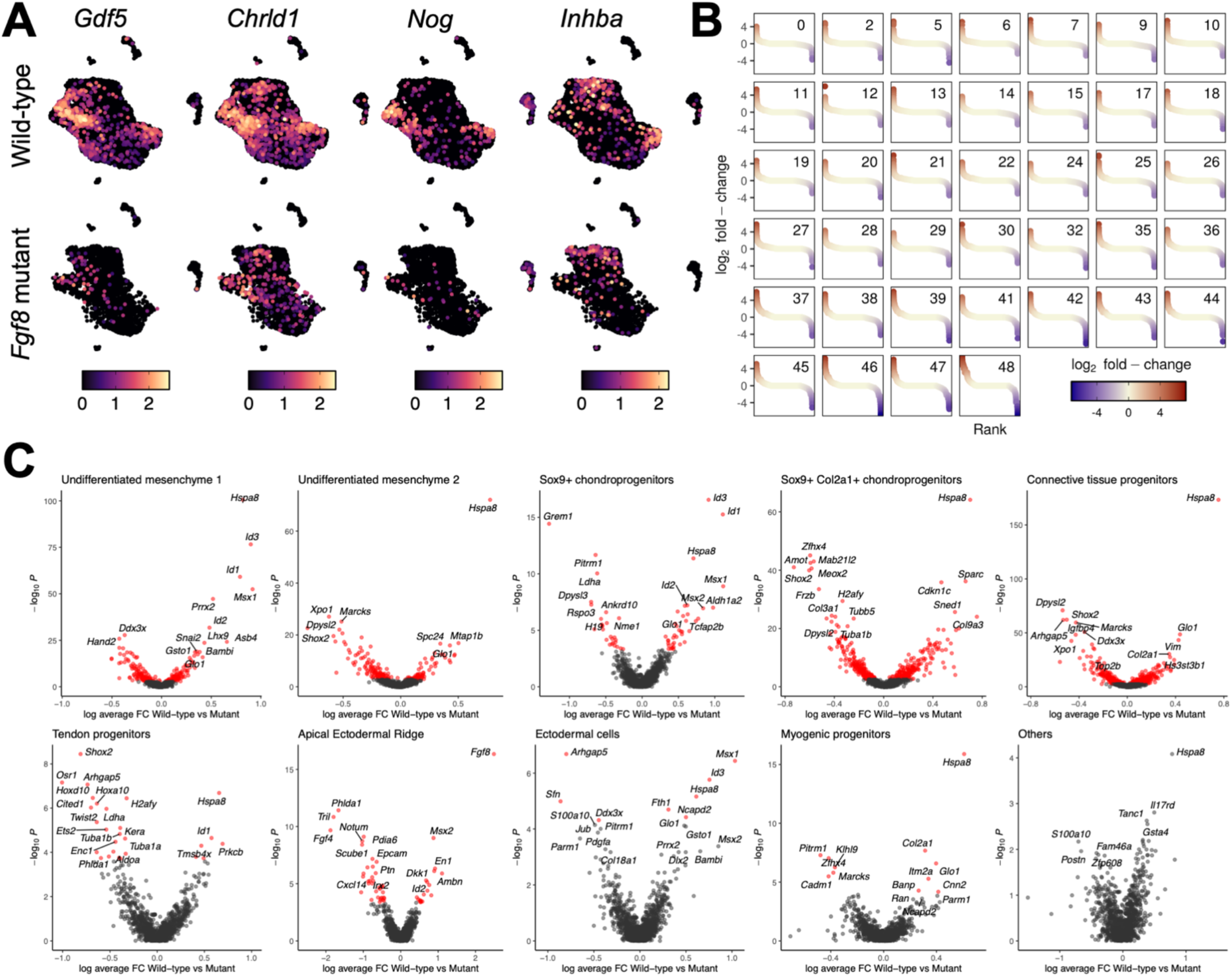
Transcriptional differences between wild-type and mutant limb cells. **(A)** UMAP showing log_10_ depth-normalized counts for some genes associated with the TGFβ superfamily signaling showing a reduction of expression in mutant cells. The data is split by genotype: wild-type (top row) and *Fgf8* mutant (bottom row). **(B)** Plot showing the distribution of log_2_ fold-changes between wild-type and mutant cells within each Seurat cluster. **(C)** Volcano plot showing the p-values of statistical tests for differential expression as a function of log_2_ fold-change between wild-type and mutant cells in each cell type. Genes for which BH-corrected p-value < 0.01 are represented in red and genes names are indicated for the top 20 most significant genes.

**Fig. S13.**
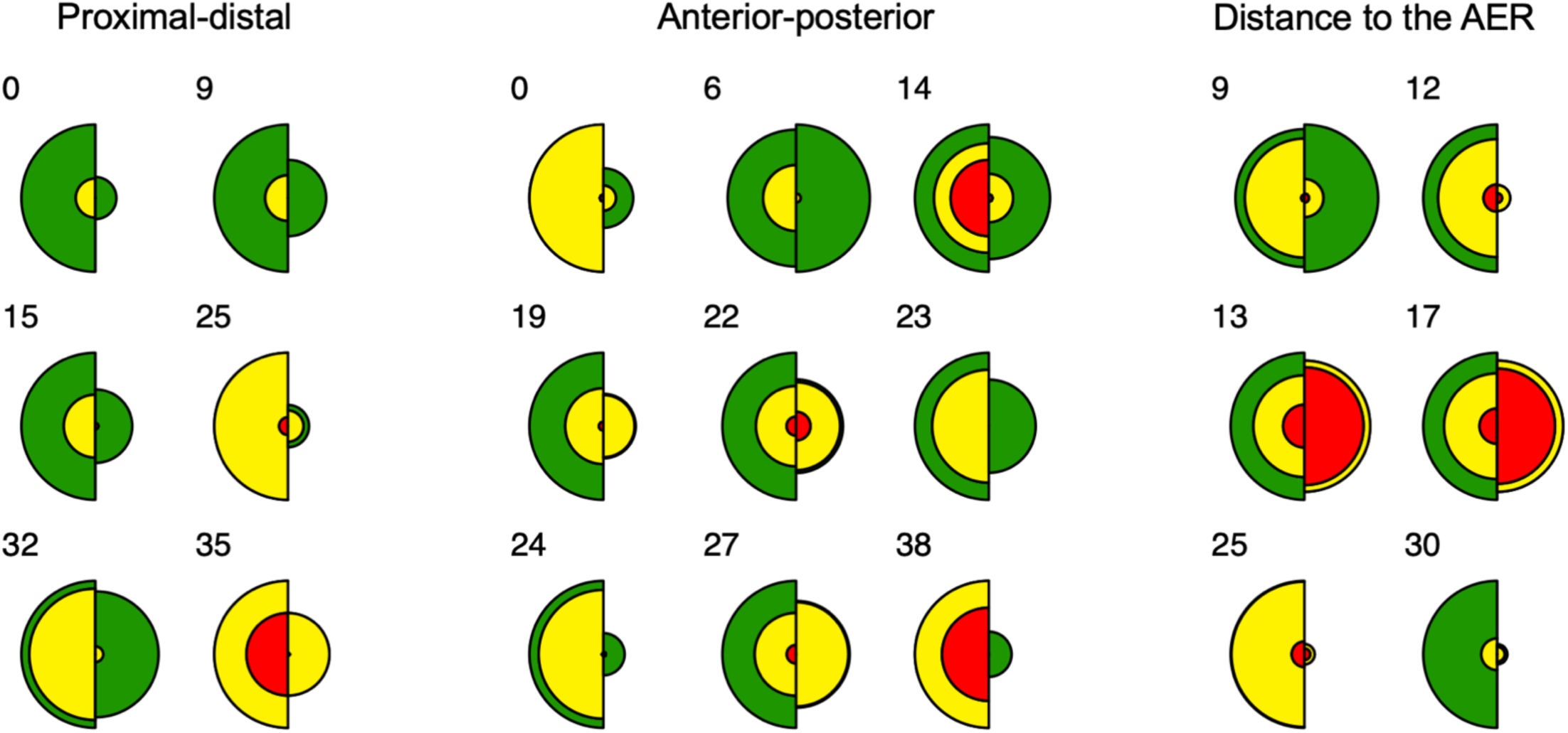
Spatial distribution changes between wild-type and mutant limb cells. Circular histograms representing the distribution of colors for wild-type and mutant cells in Seurat clusters that show altered spatial distributions.

